# Phenotypic evolutionary response to temporally limited pollinator access in Brassica rapa

**DOI:** 10.64898/2026.01.23.701299

**Authors:** Authier Elisabeth, Aeschbacher Simon, Frachon Léa

## Abstract

Plant–pollinator interactions are essential for plant reproductive success and pollinator food supply. However, the ongoing pollinator decline threatens many wild and cultivated flowering plants, urgently requiring studies on its impact on plant populations and their potential evolutionary responses. We combined an experimental evolution with a resurrection approach to test phenotypic evolutive changes in response to artificially limited access of plants to natural pollinators in a common garden setting using *Brassica rapa*. After six generations, we detected putative adaptive responses to the pollination treatments, including changes in phenology, floral morphology and flower volatile organic compounds, associated with changes in overall attractiveness to hoverflies. Although the generalist plant B. rapa shows ability to rapidly respond to strong limitations to its natural pollinator community, the observed decrease of fitness could threaten the population.

## Introduction

About 88% of all species of flowering plants worldwide are pollinated by animal pollinators (Ollerton *et al*. 2011) and depend on them for their reproduction. However, in the last few decades, a worldwide decline of domesticated and wild insect pollinators has been observed (Kluser & Peduzzi 2007; Potts *et al*. 2010). This loss of pollinators has also been observed to negatively impact ecosystem services and food production (Aizen *et al*. 2009; Klein *et al*. 2007; Singh & Adhikary 2021). For instance, local declines in bee and hoverfly species abundance in Britain and the Netherlands over the past century (Biesmeijer *et al*. 2006; Powney *et al*. 2019) support growing concerns about the persistence of local pollinator communities. Indeed, a decrease in pollinator diversity and abundance leads to pollen limitation and reduces plant reproductive success (Bennett *et al*. 2020; Thomann *et al*. 2013). For instance, lower insect pollinator diversity negatively affects fruit and seed set in *Raphanus sativus* (Albrecht *et al*. 2012).

The extent to which a decrease in pollinator diversity and abundance affects plants may depend on the degree to which plants are specialised to their pollinators (Ramos-Jiliberto *et al*. 2020). Generalist-pollinated plant species, *i.e.*, plant species interacting with a wide diversity of pollinator species for their reproductive success (Ollerton *et al*. 2007), play a central role in plant–pollinator networks due to their higher connectivity than other specialist-pollinated plant species (Bascompte & Jordano 2007). Indeed, thanks to their broader interaction networks and inherent resilience, generalist plants are less prone to extinction and potentially better able to adapt to pollinator loss than specialist species. However, few studies have examined the adaptive potential of these generalist-pollinated plant species in response to pollinator declines or to the destabilization of their pollination networks. Understanding the adaptive potential of generalist plants is particularly critical in crops, where pollinator declines could directly threaten crop yields and food security (Klein *et al*. 2007, Aizen *et al*. 2009). Moreover, the loss of generalist plant species can destabilize plant–pollinator interactions, decrease pollinator abundance (Biella *et al*. 2019; Palacio *et al*. 2016), and lead to chain extinctions of plant and pollinator species from the ecological network (Bascompte *et al*. 2019). Understanding whether and how generalist-pollinated species respond to changes in access to pollinators is crucial for ensuring ecosystem stability. However, only a few studies have explored the adaptive response of generalist plant species to pollinator community disturbance. For instance, rapid phenotypic responses to a single pollinator have been observed in generalists *Mimulus guttatus* (Bodbyl Roels & Kelly 2011) and *Brassica rapa* (Gervasi & Schiestl 2017) in greenhouse conditions. Interestingly, Schiestl *et al*. (2018) found that *Brassica rapa* exposed to a synthetic pollinator community with one bumblebee and one hoverfly species under controlled conditions evolved a ‘generalised-pollination phenotype’ overlapping with, but distinct from, the phenotypes seen under pollination by only one of the two pollinator species. However, in natural ecosystems, generalist plants interact with complex pollinator communities involving more than two pollinator species. To our knowledge, the evolutionary response of generalist plant species to changes in the extent of access to their natural pollinator communities under field conditions remains unknown.

In response to limited access to pollinators, flowering plant species can adopt two opposite strategies: decreasing their dependency on pollinators or reinforcing their interaction with pollinators (Thomann *et al*. 2013). Specifically, the first strategy for plants in response to limited access to pollinators is to enhance selfing ability, *i.e.*, reproducing using their own pollen, ensuring reproductive success in the absence of pollinators (Lloyd 1992) and is referred to as reproductive assurance. By default, selfing is suppressed in many plant species by physical and molecular barriers (Rea & Nasrallah 2008; Wang & Filatov 2023). However, these barriers can disappear within just a few generations in the absence of pollinators, as demonstrated by increased selfing ability in *Mimulus guttatus* experimental evolution study (Bodbyl Roels & Kelly 2011). Similarly, selfing ability increased in *Brassica rapa* in response to less efficient pollinators for pollination, like hoverflies (Gervasi & Schiestl 2017). In these two examples, the increase in selfing ability was associated with a decrease in herkogamy, a key physical barrier defined as the distance between the stigma and anthers in individual flowers, which promotes gamete contact within the same flower (Opedal 2018). However, the evolution of selfing is complex and involves a set of distinct traits, collectively termed the ‘selfing syndrome’ such as smaller flowers, decreased scent emission, *etc.* (Barrett 2003; Cutter 2019). However, this increased selfing capacity also leads to population-level costs, such as reduced genetic diversity. A second potential strategy in response to limited access to pollinators is to strengthen interactions with pollinators (Thomann *et al*. 2013) by increasing plant attractiveness (pollinator visitation rates) and pollen transfer efficiency. Flower traits that determine flower attractiveness are associated with pollinator detection and access to flowers (Bauer *et al*. 2017; Gervasi & Schiestl 2017) such as plant size, flower number, flower morphology, flower colour, composition and amount of nectar, and floral scent produced (Klinkhamer & De Jong 1993; Klumpers *et al*. 2019; Majetic *et al*. 2009; Wright & Schiestl 2009). However, it remains unclear whether plants preferentially adopt one strategy over the other in response to limited pollinator access.

An efficient method to study plant responses to environmental changes is an experimental evolution study (Kawecki *et al*. 2012). Experimental evolution offers the possibility of manipulating a set of environmental conditions to exert selection pressures (Bennett & Lenski 1999). However, if the aim of an experimental evolution study is to quantify the evolutionary response of an organism to a set of manipulated conditions under otherwise natural conditions, it may be difficult to separate the effect of the focal conditions from those of the uncontrolled natural conditions. Thus, such an in situ experimental evolution approach provides ecological realism but also introduces the possibility of confounding environmental effects. For instance, climatic conditions cannot be fully controlled in *in situ* experiments but may elicit responses that enhance or attenuate the effects of the experimental manipulation. Distinguishing the effects of the experimental manipulation from those of the climate therefore requires recording the climatic conditions over the course of the experiment and including them as explanatory covariates in the statistical analyses. Moreover, the expression of genetically determined focal traits may be affected by uncontrolled environmental conditions and parental effects. These factors may thus confound the comparison of traits measured before and after the experimental treatment. In plants, it is possible to mitigate both of these factors by regenerating from stored seeds the generations from before and after the evolution experiment under the same controlled conditions in a so-called resurrection approach (Franks *et al*. 2008, 2018; Weider *et al*. 2017).

The overarching aim of our study is to better understand how generalist-pollinated flowering plant species respond to temporal limitation in access to their natural pollinator community. To achieve this aim, we designed an experimental evolution study to assess i) whether and how morphological traits and fitness components respond, and ii) how the attractiveness of plants to pollinators evolves.

## Material and Methods

### Study system and experimental evolution design

We used the fast-cycling standard variety of *Brassica rapa* from Wisconsin Fast Plants® (Carolina Biological Supply, Burlington, USA) as our study system (Wendell & Pickard 2007). We randomly divided 108 full-sib seed families (supplementary information) into three sets of 36, hereafter referred to as “replicates”. Each replicate underwent an independent six-generation evolutionary experiment with three pollination treatments (“Pollination treatments” subsection). Throughout the manuscript, we refer to a treatment × replicate combination as a “replicate population”. We started the experiment with nine initial replicate populations (3 treatments × 3 replicates) and propagated these nine replicate populations individually over the experimental generations for each of the three pollination treatments (Fig. 1, supplementary information).

**FIGURE 1.**
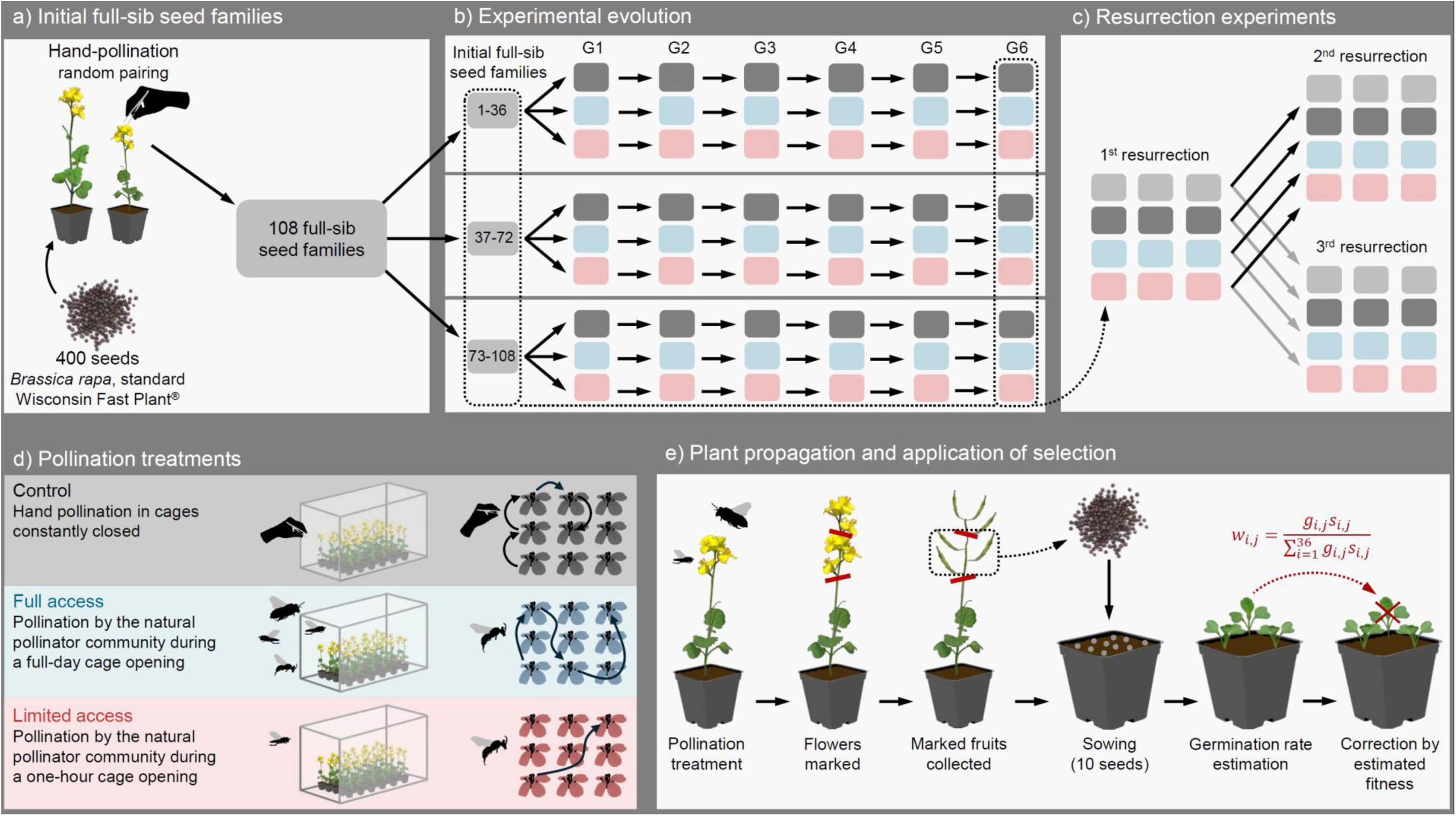
Experimental design. a) 108 full sib-seed families of fast-cycling B. Rapa have been generated from 400 B. rapa seeds, and b) propagated over six generations (G1 to G6) in three pollination treatments in common garden, each comprising three parallel replicate populations of 36 individuals. c) To remove maternal and environmental effects, seeds from the full-sib families and from the sixth generations of the experimental evolution were regrown in a resurrection experiment under similar greenhouse conditions for two generations (first and second resurrections). Due to set up constraint, we regrow a second time the seeds of the first resurrection, leading to a third resurrection. d) Treatments included one control treatment and two pollination treatments in which access of pollinators was temporally limited. e) To propagate plants from one generation to the next, we estimated relative fitness from the product of the total number of seeds produced times the germination rate and corrected the number of seedlings in each pot by the estimated relative fitness of its maternal plant.

We grew the 324 individual plants (9 replicate populations x 36 individual plants) in a phytotron for two weeks, potted them in standardized soil, and moved them into nine outdoor cages surrounded by an insect-proof net that were set up at the Botanical Garden of the University of Zürich (Fig. S1). Inside each cage, we randomly positioned 36 individual plants from a given replicate population (Fig. S1).

During the experimental evolution study, we measured 17 traits including seven architectural and morphological traits, and 10 fitness-related traits at each generation (see table S1, supplementary information). Climate data was recorded (temperature and light intensity, supplementary information) using a datalogger installed in the central outdoor cage, and principal component analysis was performed to summarize the variation of the 10 climatic variables (Table S2).

### Pollination treatments

After moving plants into outdoor cages, we applied three pollination treatments by artificially manipulating temporal pollinator access to simulate a decrease in pollinator richness and abundance. 1) Control treatment: we hand-pollinated plants by transferring pollen from flowers of a given plant to one to six flowers of the nearest plant. We kept the three cages assigned to the three replicates in this treatment constantly closed to exclude insect pollinators. 2) Full Access treatment: we let the natural surrounding insect pollinator community pollinate the plants. We opened each of the three cages once during the life cycle for five consecutive hours during the highest pollinator activity (between 11 am and 4 pm). 3) Limited Access treatment: we temporally reduced the access of the natural pollinator community to plants compared to the Full-Access treatment. We opened each of the three cages assigned to the three replicates in this treatment once during the life cycle for one hour during the highest pollinator activity (between 12 pm and 1 pm). Our observations of pollinator visits during treatment application (supplementary information) revealed significantly lower pollinator abundance and diversity in the Limited Access treatment compared to the Full Access one (Supplementary information, Table S3, Dataset 1).

### Resurrection experiments

To reduce maternal and environmental effects that could confound generation effects, we conducted two refresher generation resurrection experiments (Fig. 1c, supplementary information). We sowed seeds from the first and sixth generations together to generate a total of 12 replicate resurrection populations (three initial replicate populations plus three treatments × three replicate populations from the sixth generation). We grew these replicate resurrection populations in a phytotron for two weeks, and then transferred them into a greenhouse (Irchel Campus, University of Zürich). Two weeks later, within each replicate resurrection population, we cross-pollinated eight flowers per plant with pooled pollen from the same replicate resurrection population. At fruit maturity, we generated a second resurrection generation to decrease maternal effect by growing one offspring per maternal plant (n = 36) following the same protocol and conditions as previously described. Finally, we repeated a third resurrection experiment in order to estimate changes in mating system using seeds from the first resurrection (Fig. 1c). In the manuscript, unless contraindicated, we use the term ‘resurrection experiment’ without distinction, referring to either the second or third resurrection depending on the context.

During the resurrection experiments, we measured 33 traits including: flowering time, eight architectural and morphological traits, 13 flower volatile organic compounds (VOCs), and 11 fitness-related traits (Table S1, supplementary information).

### Pollinator preference

To investigate the effect of pollination treatments on the overall flower attractiveness to pollinators, we performed a four-choice test. In this test, an insect pollinator was given a choice among four focal plant individuals from the four different resurrection populations (initial generation, sixth generation of Control, Full Access, and Limited Access). We used the pollinator’s first choice as a preference proxy. During the four-choice test, we released one bumblebee (*Bombus terrestris*) or one adult hoverfly (*Episyrphus balteatus*) inside a cage. In total, we performed 106 four-choice tests for each pollinator species. On the test day, we counted the *number of open flowers* for each of the four focal plants.

### Statistical analyses

We tracked phenotypic changes over time during the experimental evolution. Based on the mean values of traits across plant individuals within each replicate population times generation, we tested separately (1) the overall effect of pollination treatment, and (2) the overall effect of generation with Kruskal—Wallis test.

To quantify both the effect of climate and that of generations, we performed two independent multi-way ANOVAs to explain trait changes by treatment, and either by generation or by climate for the ten traits showing a normal distribution in R v4.4.1 according to the following equations:

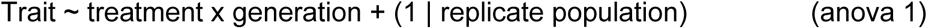

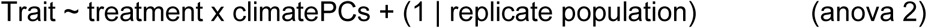

where *Trait* represents the mean of 10 phenotypic and fitness-related traits, *treatment* the three pollination treatments, *generation* a categorial explanatory variable representing the six generations, *climatePCs* the first two principal components derived from the climate PCA, and *replicate population* as random factor.

To detect phenotypic and fitness-related trait evolutionary changes, we used arithmetic mean for each trait from the resurrection experiment dataset. First, to understand changes in trait relationships across treatments, we calculated Spearman’s pairwise rank correlation coefficients (package Hmisc v5.0-1, Harrell Jr 2003) in R environment among traits within each four resurrection populations. Additionally, to assess evolutionary changes occurred over six generations, we performed Kruskal—Wallis tests on arithmetic mean of 33 traits within each replicate resurrection population, with treatment as a categorical explanatory variable. For traits with significant results (p-value < 0.05), we performed multiple pairwise Kruskal-Wallis tests (R package pgirmess v 2.0.3, Giraudoux *et al*. 2024).

To quantify directional selection, we applied the Lande & Arnold framework (Lande & Arnold 1983) to explain the effect of uncorrelated (significant rho_Spearman_ < 0.8) phenotypic traits (fixed-effect explanatory variables) on relative fitness estimate. We performed three directional selection analysis based (1) on five uncorrelated phenotypic traits that passed the Shapiro test, (2) on seven uncorrelated phenotypic traits related to phenology and floral morphology traits, and (3) on 11 uncorrelated traits associated with VOCs. The relative fitness estimate was the relative number of seeds per plant within the respective replicate resurrection population. We included replicate as a random slope, and the phenotypic traits were centred and scaled within their replicate resurrection population for each generation. We interpreted the partial regression coefficients pertaining to the explanatory variables as directional selection gradients.

To test changes in overall plant attractiveness, we performed a four-choice experiment and tested the pollinator preference for one of the four resurrection treatments. We performed pairwise comparison of proportions (pairwise.prop.test function) of chosen plants between the initial and the sixth generation for the three treatments. To see if the distribution of the phenotypic traits differs between plants chosen and not chosen by pollinators, we performed pairwise Wilcoxon tests in R for each phenological and phenotypic trait by using the arithmetic means between plants chosen and not chosen by either bumblebees or hoverflies.

## Results

### Tracking phenotypic changes over generations

We assessed the overall effects of treatment and generation on 17 traits using pairwise Kruskal-Wallis tests. Additionally, to disentangle the potential confounding effects between generation and climate, we performed two multi-way ANOVAs: included treatment and generation as factors (anova 1), or treatment and climate as factors (anova 2). First, we observed a significant overall effect of treatment on seven out of ten fitness-related traits, but no treatment effect on floral morphological traits (Table S4). Second, we observed a significant overall effect of generation (considering all replicate population together) on 11 traits related to morphological traits and fitness-related traits (Fig. 2, Table S4). We also detected significant effects of both generation (anova 1) and climate (anova 2) for all tested traits (“within” replicate populations in Dataset 2), as illustrated by mean flower diameter (Fig. 2a). These results revealed variations across generations, which is confounded by the distinct climate experienced by each generation, emphasizing the importance of combining experimental evolution in natural condition with resurrection approach.

**FIGURE 2.**
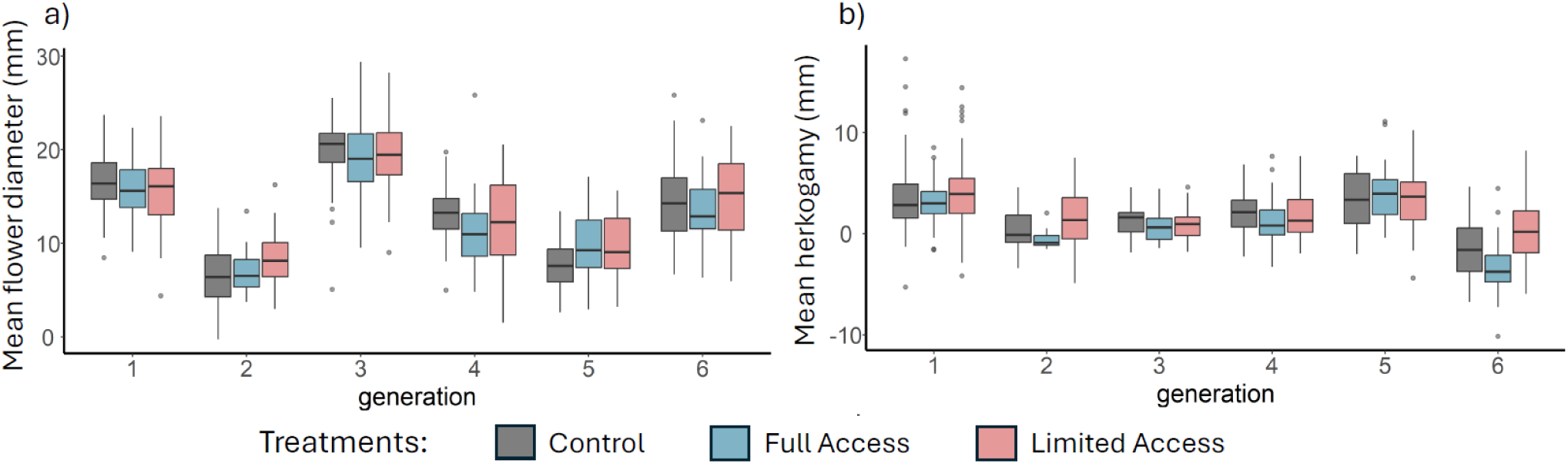
Tracking phenotypic trait evolution across six generations of the experimental evolution study in common garden. Empirical distribution of (a) mean flower diameter and (b) mean herkogamy per pollination treatment as measured directly in the experimental evolution study.

### Evolutionary changes in phenotypic trait correlations

To assess the variation in phenotypic correlations between generations and according to treatments, pairwise Spearman correlations among 30 phenotypic traits and fitness-related traits measured in resurrection experiment were performed. These correlations showed different patterns according to the resurrection treatments (initial generation, three evolved pollination treatments), and to the trait sets considered (Fig. S2). For instance, we observed a decrease in the strength of positive correlations between the flowering time and flower morphology traits from the initial generation to the last generation across all treatments (Fig. S2). Moreover, we observed a shift from negative correlation in initial generation to positive correlation between some flower morphology traits and amounts of flower scent components, especially in Full and Limited access treatments (Fig. S2). In contrast, a slight change in correlation from positive to negative appeared between fitness-related traits and the floral VOCs from initial to last generation for Control and Limited access treatments (Fig. S2). Finally, for certain sets of traits, the correlations within them remain overall stable during the evolutionary process like the floral VOCs, the floral morphological traits, and the fitness-related traits.

### Rapid phenotypic evolutionary response to pollination treatments

To unravel the phenotypic differentiation among the replicate resurrection populations, we performed a linear discriminant analysis (LDA) on different categories of traits. We observed an overall phenotypic differentiation among the initial generation and the replicate resurrection populations of the last generation (Fig. S3). We found phenotypic differentiation among treatment populations in LDA based on floral morphological traits and on fitness-related traits (Fig. S3ac) highlighted by a non-overlapping of centroid. However, these phenotypic differentiation within trait categories are slight, as indicated by the overlap of the ellipses in the space described in LDA. In contrast, in LDA based on floral VOCs, the centroid of replicate populations from evolved pollination treatments clustered together but remained distinct from the initial generation (Fig. S3e). These results indicated some phenotypic changes of combination of traits in response to different selection-mediated treatments.

To investigate the treatment effect (initial generation and the three evolved pollination treatments) on phenotypic trait variations, we performed Kruskal—Wallis test. Overall, we observed significant evolutionary changes across treatments for 19 out of 33 measured traits related to morphological and fitness-related traits i.e. significant differences between the initial generation and at least one of the evolved populations (Table 1). Interestingly, our findings revealed significant differences between the initial and the last generations in the Full and/or

**TABLE 1.** Results of Kruskal—Wallis tests to explain the variation of 33 traits measured in the resurrection experiments by treatment as a categorial explanatory variable. Two sets of tests have been performed: (1) an overall Kruskal—Wallis tests and (2) multiple pairwise Kruskal—Wallis tests (between each per of treatments). For each trait, the arithmetic mean within each treatment has been calculated and indicated on the right of the table. The four pollination treatments are: Initial sib-seed families ("G0"), Control ("C"), Full access ("FA"), and Limited access ("LA"). Significant p-values (<0.05) are indicated in bold.

| Traits | Kruskal—Wallis test |  | pairwise Kruskal—Wallis test |  |  |  |  |  | Mean |  |  |  |
| --- | --- | --- | --- | --- | --- | --- | --- | --- | --- | --- | --- | --- |
|  | chi-squared | p-values | G0 - C | G0 - FA | G0 - LA | C - FA | C - LA | FA - LA | G0 | C | FA | LA |
| Flowering time | **58.8** | **<0.001** |  | Yes | Yes | Yes | Yes |  | **19.71** | **20.22** | **20.88** | **21.23** |
| Height of the plant | **12.6** | **0.006** |  |  |  | Yes | Yes |  | 30.01 | **30.79** | **28.61** | **28.75** |
| Number of inflorescences | **21.1** | **<0.001** |  |  |  | Yes | Yes |  | 1.61 | **1.43** | **1.83** | **1.76** |
| Mean flower diameter | 4.4 | 0.220 |  |  |  |  |  |  | 13.03 | 13.53 | 13.08 | 13.44 |
| Mean petal length | **10.7** | **0.014** |  |  | Yes |  |  |  | **9.68** | 9.23 | 9.02 | **9.00** |
| Mean petal width | **13.7** | **0.003** |  |  | Yes |  |  |  | **4.01** | 3.62 | 3.75 | **3.49** |
| Mean pistil length | 3.1 | 0.379 |  |  |  |  |  |  | 6.33 | 6.06 | 5.60 | 5.20 |
| Mean stamen length | 3.2 | 0.366 |  |  |  |  |  |  | 5.93 | 5.86 | 5.88 | 5.71 |
| Mean herkogamy | 2.4 | 0.489 |  |  |  |  |  |  | 0.40 | 0.20 | -0.28 | -0.51 |
| Total number of flowers | **11.4** | **0.010** |  | Yes |  |  |  | Yes | **17.56** | 18.31 | **19.16** | **18.24** |
| Total number of seeds | 7.5 | 0.058 |  |  |  |  |  |  | 63.32 | 60.25 | 53.89 | 54.55 |
| Total seed weight | **40.9** | **<0.001** | Yes | Yes |  | Yes |  | Yes | **90.29** | **74.32** | **59.25** | **77.64** |
| Mean seed weight | **55.6** | **<0.001** | Yes | Yes |  | Yes | Yes | Yes | **1.53** | **1.32** | **1.16** | **1.59** |
| Total seed weight per fruit | **18.7** | **<0.001** | Yes | Yes | Yes |  |  |  | **27.43** | **22.85** | **21.87** | **22.04** |
| Benzaldehyde | **11.1** | **0.011** |  |  | Yes |  |  |  | **2.07** | 1.16 | 1.43 | **1.09** |
| β-Pinene | 4.8 | 0.184 |  |  |  |  |  |  | 0.28 | 0.20 | 0.10 | 0.06 |
| 1-Butene-4-isothiocyanate | **48.0** | **<0.001** | Yes | Yes | Yes |  |  | Yes | **4.26** | **2.15** | **1.95** | **1.47** |
| Z-(3)-Hexenyl acetate | **26.2** | **<0.001** | Yes | Yes | Yes |  |  |  | **0.44** | **0.29** | **0.21** | **0.18** |
| Limonene | **59.5** | **<0.001** | Yes | Yes | Yes |  |  | Yes | **4.41** | **2.34** | **2.19** | **1.44** |
| Methyl benzoate | **27.7** | **<0.001** |  |  | Yes |  |  | Yes | **0.14** | 0.21 | **0.16** | **0.27** |
| Nonanal | **35.1** | **<0.001** |  | Yes | Yes |  | Yes |  | **2.62** | **2.07** | **1.60** | **1.36** |
| Phenylethyl alcohol | 2.8 | 0.416 |  |  |  |  |  |  | 0.09 | 0.07 | 0.04 | 0.05 |
| Benzyl nitrile | **61.0** | **<0.001** |  | Yes | Yes |  | Yes | Yes | **0.88** | **0.84** | **0.59** | **0.31** |
| Methyl salicylate | **13.9** | **0.003** |  |  | Yes |  |  |  | **0.07** | 0.06 | 0.05 | **0.04** |
| 2-Amino benzaldehyde | **28.0** | **<0.001** |  | Yes | Yes |  |  |  | **0.16** | 0.13 | **0.08** | **0.09** |
| p-Anisaldehyde | **25.4** | **<0.001** |  | Yes | Yes | Yes | Yes |  | **0.10** | **0.09** | **0.07** | **0.05** |
| Methyl anthranilate | **20.0** | **<0.001** |  |  |  | Yes |  | Yes | 0.22 | **0.07** | **0.01** | **0.08** |
| mean fruit length under outcrossing | **48.1** | **<0.001** |  | Yes | Yes |  | Yes | Yes | **55.78** | **53.83** | **49.25** | **43.31** |
| mean fruit length under autogamy | **9.9** | **0.020** |  |  |  |  |  | Yes | 41.29 | 42.67 | **45.87** | **38.31** |
| mean nb of seeds per fruit under outcrossing | **24.3** | **<0.001** |  | Yes | Yes |  | Yes |  | **16.94** | **14.24** | **13.44** | **10.80** |
| mean nb of seeds per fruit under autogamy | **10.5** | **0.015** |  |  |  | Yes |  |  | 6.19 | **4.96** | **9.37** | 6.37 |
| number of seeds/mean fruit length under outcrossing | 7.3 | 0.064 |  |  |  |  |  |  | 0.30 | 0.25 | 0.27 | 0.24 |
| number of seeds/mean fruit length under autogamy | **9.1** | **0.028** |  |  |  |  |  |  | 0.17 | 0.14 | 0.21 | 0.19 |

Limited Access treatment, but not in the Control, across 13 traits associated with different trait categories, highlighting evolutionary response to the pollination treatment, rather than uncontrolled environmental effects that would also be present in the control treatment. For instance, we detected a significant delay in flowering time for both Limited and Full access treatments, but not in Control treatment (Fig. 3a, Table 1). Moreover, we observed significant evolutionary changes for 11 out of 13 floral VOCs with, for instance, a decrease in benzyl nitrile in Limited Access and Full Access compared to the initial generation (Fig. 3b, Table 1). Finally, we detected significant evolutionary changes in outcrossing ability with a decrease in fruit length and seed weight across generations in Limited and Full Access treatments (Fig. 3c, Table 1). However, we did not observe significant evolutionary changes for autogamy ability for none of the pollination treatments (Table 1). Finally, we observed evolutionary changes in six traits where significant changes occurred in the Limited and/or Full Access treatments, but also in the Control treatment (Table 1). This introduces uncertainty in attributing these evolutionary changes to the pollination treatment(s) rather than uncontrolled environmental effects selecting in our populations.

**FIGURE 3.**
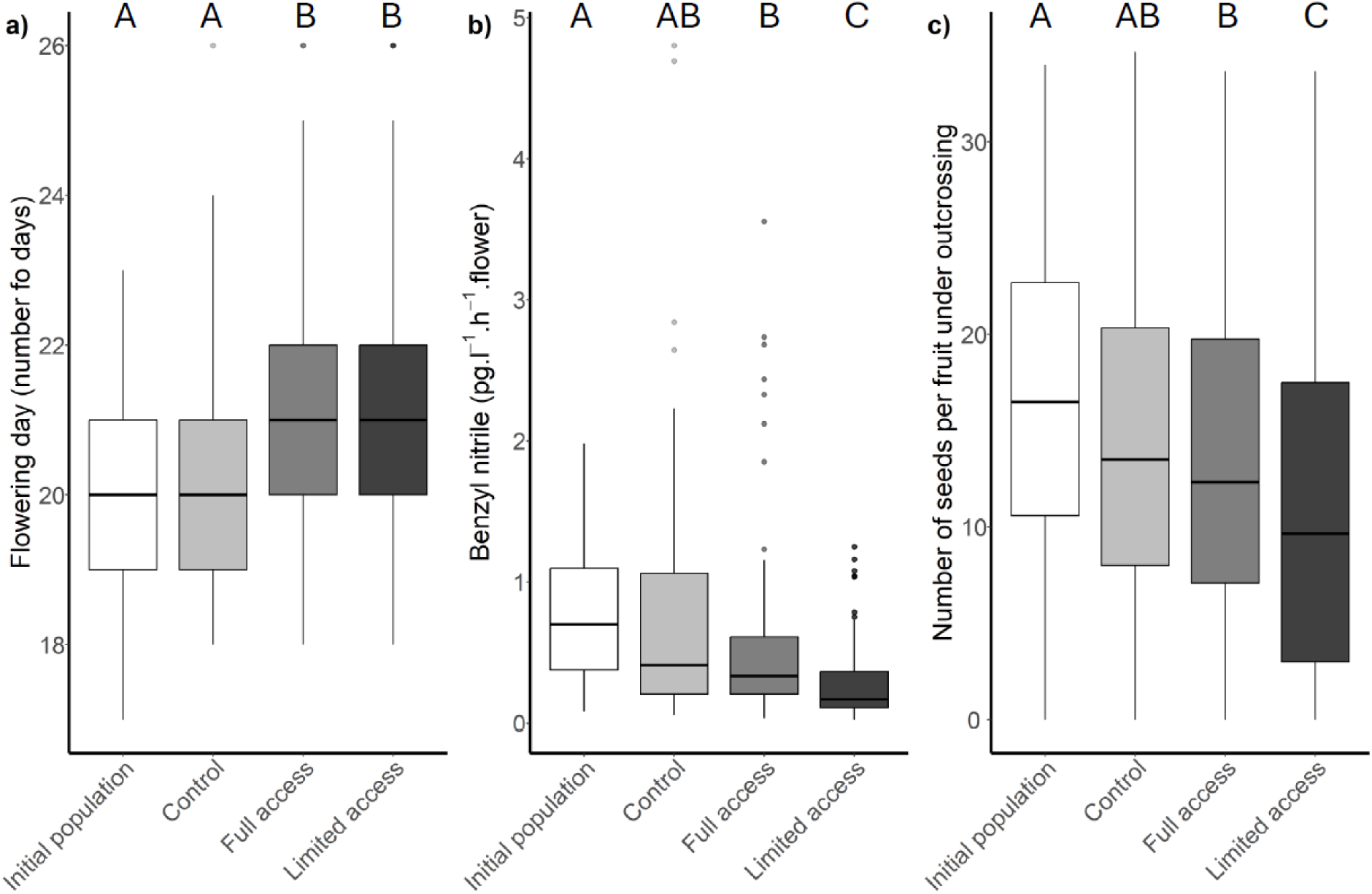
Evolutionary changes observed in resurrection approach for the different pollination treatments. Empirical distribution of a) the flowering time; b) the amount of benzyl nitrile in floral VOCs; and (c) the number of seeds per fruit under outcrossing. For each plot, the initial populations and the three evolved populations (Control, Full Access, and Limited Access) are represented. Significant pairwise comparison between means of distributions (pairwise Kruskal-Wallis test) are emphasized.

### Changes in the strength of directional selection

Finally, to quantify changes in the strength and the direction of the selection for phenotypic traits before and after potential selection within different pollination treatments, we investigated directional selection analyses in resurrection experiment. These analyses have been performed on two different set of traits: (1) with all plant architecture and flower morphology; and (2) VOCs floral traits. In initial generation, two phenotypic traits were under positive selection (height of the plant, number of inflorescences, Table S5a), and one under negative selection (pistil length). The strength of directional selection increased in the control treatment for these traits. However, we observed different patterns for both pollination treatments. In the Full Access, we observed the emergence of negative selection for flowering time, and positive selection for petal width (Table S5a). In the Limited Access, we observed a strengthening of positive selection on plant height (Table S5a). Finally, we did not observe directional selection in traits related to scent compounds, either in the initial generation or in evolved populations (Table S5b).

### Plant evolution driven by pollinator preferences

To determine whether the phenotypic responses observed in our experiment is associated with evolutionary changes in overall plant attractiveness, we tested for potential shifts in pollinator preference among resurrection treatments (initial generation, Control, Limited Access, and Full Access). In four-choice test experiment, we assessed the preference of two common pollinators (bumblebees and hoverflies) in our common garden that visited our plants during experimental evolution (Dataset 1). We did not observe distinct pollinator preferences between the initial generation and the evolved populations, except for a lower preference of hoverflies for plants in Limited Access over the ones from initial generation (Fig. 4ad, Table S6).

Moreover, we tested the overall preference of bumblebee and hoverfly for specific phenotypic traits. For instance, we observed a preference for high number of flowers for both species of pollinators (Fig. 4b, Table S6), as well as a preference of bumblebees for earlier flowering time (Table S6). Additionally, we observed a significant preference of bumblebees for smaller pistil length, negative herkogamy, and lower amount of methyl benzoate and higher amount of phenylethyl alcohol in floral VOCs (Fig. 4, Table S6). Similarly, we found that hoverflies preferred higher amount of limonene and methyl salicylate, and lower amount of 2-amino benzaldehyde (Table S6). Lastly, we observed a shared preference of bumblebees and hoverflies for higher amount of nonanal and benzyl nitrile in floral VOCs (Fig. 4e and f, Table S6).

**FIGURE 4.**
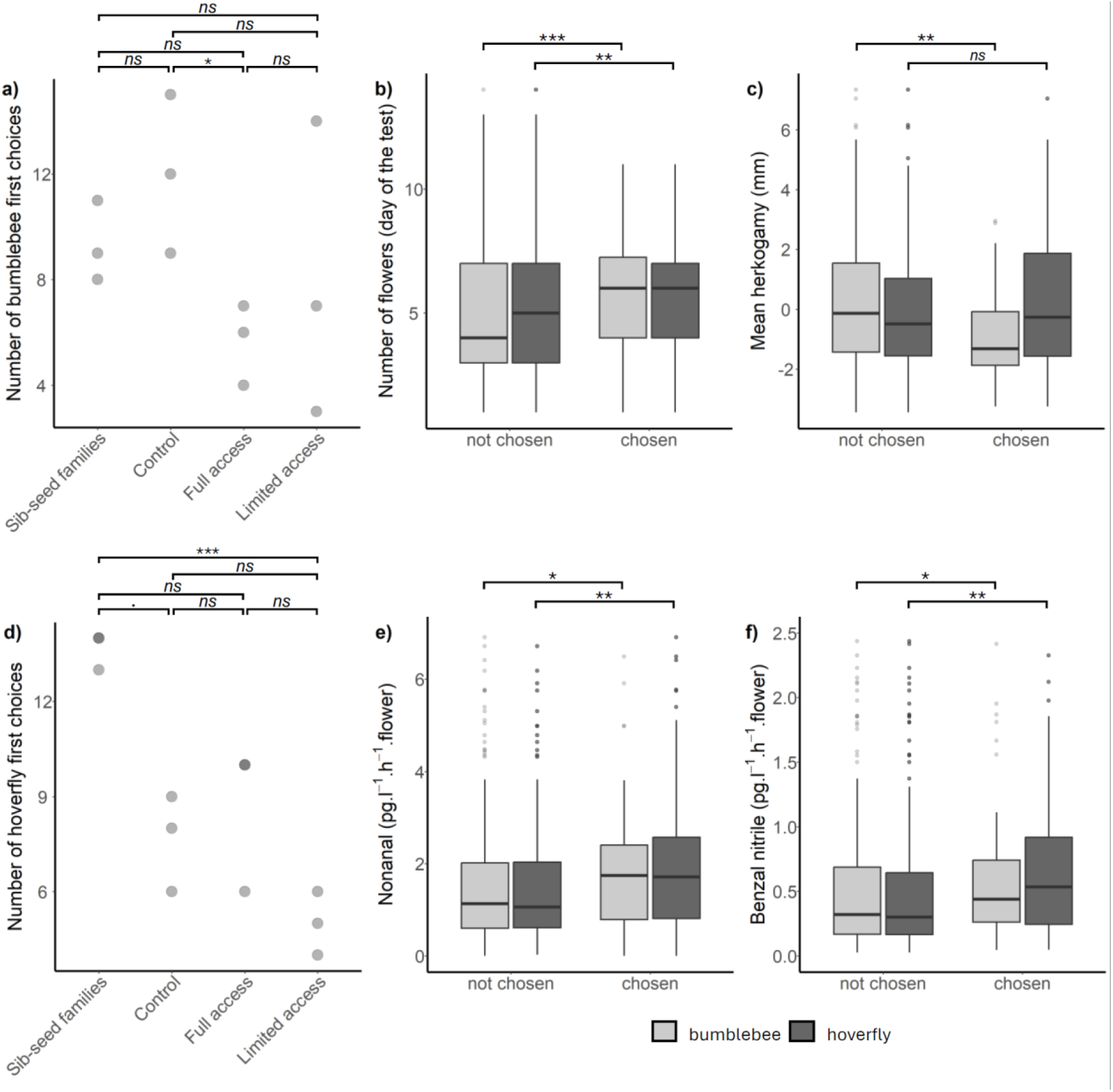
Preference of Bombus terrestris and Episyrphus balteatus for treatment and phenotypic traits. Preference of a) bumblebees (Bombus terrestris) and d) hoverflies (Episyrphus balteatus) for resurrection pollination treatments. Distributions of trait values of plants chosen vs. not chosen as the first preference by bumblebees (light bars) and hoverflies (dark bars) for b) the number of flowers, c) herkogamy, and amounts of e) nonanal and f) benzyl nitrile in floral VOCs. The ‘number of first choices’ refers to the number of times a plant from a specific treatment was chosen as the first plant visited during the four-choice tests. For a) and b), significant pairwise comparisons of proportions are emphasized (***: 0 ≤ p ≤ 0.001, **: 0.001 < p ≤ 0.01, *: 0.01 < p ≤ 0.05, ▪: 0.05 < p < 0.1, ns: non-significant). For c), d), e), and f) significant pairwise Wilcoxon tests are emphasized (***: 0 ≤ p ≤ 0.001, **: 0.001 < p ≤ 0.01, *: 0.01 < p ≤ 0.05, ns: non-significant).

## Discussion

In response to disturbances in natural pollinator communities, generalist flowering plants may rapidly adapt by increasing their pollinator attractiveness or enhancing selfing ability. In a six-generation common-garden evolutionary experiment with three varying temporal access to natural pollinators, we observed evolutionary changes in morphological and fitness-related traits. However, distinguishing these changes from the effects of distinct environmental conditions across generations is challenging. To minimize environmental and maternal effects, we conducted a resurrection experiment in a controlled greenhouse, growing together the initial and the last generations. We found rapid phenotypic changes in response to variation in pollinator access, including i) reduced outcrossing ability under Limited and Full Access and ii) shifts in phenology (delay of flowering), floral morphology (decrease in petal size), and floral scent under Limited Access. Finally, contrary to our assumptions, we detected reduced attractiveness to hoverflies in Limited Access.

Our resurrection approach revealed that temporally limited access to natural pollinators can drive rapid shifts in plant mating systems. Specifically, we observed a decrease in outcrossing ability in both the Limited and Full Access treatments. This mating system shift was associated with reduced fruit length under outcrossing, and fewer seeds per fruit under outcrossing, with higher effect in the Limited Access. Such reduction in outcrossing ability was also documented in a controlled greenhouse evolutionary experiment in response to hoverfly pollination (Gervasi and Schiestl 2017, Kofler *et al*. 2024). These changes could result from a decrease in pollinator visits, which may reduce pollen quantity — suggested but not measured — potentially leading to pollen limitation (Burd 1994). However, while a decrease in outcrossing ability is known as adaptive response to reduced pollinator availability, it is often associated with an increase in selfing (Acoca-Pidolle *et al*. 2024; Cheptou *et al*. 2022; Bodbyl Roels & Kelly 2011). In our study, we did not observe such significant changes; however, we did observe a non-significant trend toward increased selfing ability. The lack of significant changes in selfing ability may reflect known barriers preventing selfing in *Brassica* (e.g., genetic self-incompatibility, floral morphology, Brugière *et al*. 2000, Nasrallah 2017, Murase *et al*. 2020), or a latent period before any evolutionary shift in selfing ability becomes detectable. Our findings suggested that small populations facing pollinator community disturbance are at high risk of decline. In fact, the reduction in outcrossing compromises genetic diversity, while the absence of increased selfing, which could provide reproductive insurance, further reduces reproductive success, creating a dual threat to population viability.

In addition, our study highlighted rapid evolution of phenology and floral morphology in response to pollination treatments. Both Limited and Full Access treatments exhibited a delay in flowering time, with a delay more pronounced in the Limited Access. In our study, this delay in flowering time was associated with a reduced preference of both bumblebees and hoverflies. Our finding is in line with previous studies in which pollinators drive changes in flowering time (Elzinga *et al*. 2007, Xu 2023). Although flowering time is often linked to climate adaptation (Fournier-Level *et al*. 2022; Geissler *et al*. 2023; Preston & Fjellheim 2022), we detected no significant differences between the initial population and the Control treatment, confirming that changes arose from pollination treatments rather than environmental factors in our study. However, while we observed this delay in flowering time, we highlighted a significant negative selection of this phenological trait in Full Access (delay flowering time correlated with lower fitness). This finding could be explained by several hypotheses: (1) selection acting on a trait negatively correlated with flowering time, (2) a recent shift in selection that has not yet manifested in phenotypic changes, or (3) selection favouring synchrony with pollinator peak activities rather than early flowering per se. Moreover, we observed shifts in floral morphology with a decrease in petal size in the Limited Access treatment. This flower reduction pattern is often observed as a response to loss of pollinators (Tusuubira and Kelly 2024, Acoca-Pidolle *et al*. 2024). Indeed, an absence of pollinator can lead to a shift in mating-system, from outcrossing animal-pollination to selfing pollination. This transition, called ‘selfing syndrome’, is accompanied by phenotypic changes reducing the plant attractiveness to pollinators, including decrease in flower size (Tsuchimatsu and Fujii, 2022). While we did not observe significant increase in selfing ability, only a non-significant trend, this floral morphological change may represent an early evolutionary shift toward a selfing syndrome, potentially leading to increased selfing rates in subsequent generations.

Additionally, we observed evolutionary changes in many floral scent compositions in only six generations. Pollinators often show strong preferences for a specific or a combination of floral VOCs, and previous studies suggested that pollinator-mediated selection can drive changes in floral scent composition in *B. rapa* (Dorey & Schiestl 2024; Gervasi & Schiestl 2017; Ramos & Schiestl 2019). Among the observed changes in VOCs, most floral VOCs are known to influence pollinator preferences. For instance, while methyl salicylate is associated with hoverfly preference, limonene, p-anisaldehyde, 2-aminobenzaldehyde, benzaldehyde, methyl salicylate, and nonanal are linked to preferences in Hymenoptera (Dötterl & Gershenzon 2023). For example, we confirmed that plants with lower nonanal levels are less frequently chosen by both bumblebees and hoverflies. Additionally, we observed evolutionary reductions in nonanal production across six generations in both Limited and Full Access treatments, further demonstrating that our pollination treatments decrease plant attractiveness. Due to the involvement of many VOCs and their shared biosynthetic pathways (Dötterl & Gershenzon 2023), the target of selection in our study is unlikely to be a specific VOC compound, as confirmed by the lack of consistent directional selection for these traits. Instead, selection likely acts on the floral scent bouquet as a whole.

Lastly, we detected an overall decrease in plant attractiveness to hoverflies in evolved plants from the Limited Access treatment, for which we observed a significantly lower abundance of hoverflies over the six generations of selection compared to the Full Access treatment. The decrease in hoverfly attractiveness in Limited Access can be explained by changes observed in a combination of phenotypic traits, including VOCs, in response to low occurrence of these pollinators. For instance, we documented evolutionary reductions in VOCs such as limonene, nonanal, and benzyl nitrile across both Limited and Full Access treatments. Correspondingly, plants producing lower levels of these compounds were significantly less attractive to hoverflies, while these compounds are known to be involved in plant attractiveness to hoverflies (Dötterl & Gershenzon 2023). Additionally, we observed that the Limited Access treatment reduced petal size, which aligns with the overall decrease in plant attractiveness. Because producing floral scent and enhancing floral visibility to improve plant attractiveness are costly (Spigler *et al*. 2020), one plausible hypothesis for this reduced hoverfly preference for plants evolved in Limited Access is that these plants have adapted by reducing their overall attractiveness in response to reduced pollinator community. In contrast, we observed no such trend in bumblebee preferences, possibly because there was no substantial difference in bumblebee abundance between the Limited and Full Access treatments. However, we tested plant attractiveness using only two common pollinators under controlled conditions, which does not reflect the ecological realism of the environments where these plants have evolved. It would therefore be valuable to test the attractiveness of the evolved plants to a broader range of pollinators, including honeybees or small bees – groups for which we observed abundance differences between treatments - as well as within the context of a complete pollinator community. Moreover, six generations are a short time period for selective processes, and the variation of our selective pressure (pollinator community composition) across generations, may have buffered the speed of selection and likely imposed diffuse selection on traits.

Overall, our study showed that temporally limiting generalist plants’ access to their natural pollinator community can drive short-term changes in mating system and overall plant attractiveness. It therefore appeared that flowering plants have substantial adaptive capacity, particularly in traits affecting pollinator attractiveness (floral morphology and scent). However, the short time period of our experimental evolution provides only a partial view of how plants evolve in response to a decline in pollinators, and the long-term consequences of pollinator decline remain unclear. Reducing pressure on natural pollinator communities should therefore remain a priority.

## Acknowledgement

We thank Markus Meierhofer, Rayko Jonas, Daniel Schlagenhauf, Matthias Furler, Franz Huber, Laura Dällenbach for their help during experiments. We thank Florian Schiestl, Tobias Züst, Maxime Bonhomme and Carolin Kosiol for helpful discussions and inputs during the study. This work was supported by the University of Zürich and the University Research Priority Program ‘Evolution in action’.

## Supplementary Materials

Dataset 1: raw data pollinator abundance and diversity

Dataset 2: result of multi-ways ANOVAs (models 1 and 2) – experimental evolution

Dataset 3: raw data individual traits – experimental evolution

Dataset 4: raw data flower morphology – experimental evolution

Dataset 5: raw data fruit length – experimental evolution

Dataset 6: temperature and light recording – experimental evolution

Dataset 7: raw data individual traits – second resurrection

Dataset 8: raw data flower morphology – second resurrection

Dataset 9: raw data fruit length – second resurrection

Dataset 10: raw data individual traits – third resurrection

Dataset 11: raw data fruit length – third resurrection

## Author Contributions

LF and SA acquired the University Research Priority Program ‘Evolution in action” funding, conceptualized and designed the experiment, EA performed the experiment with the contribution of LF, EA performed the analyses, EA wrote the original draft, LF and SA reviewed and edited the manuscript.

## Data Availability Statement

Data and code are available in Zenodo (10.5281/zenodo.15425805).

