## Supplementary information for "Phenotypic evolutionary response to temporally limited pollinator access in Brassica rapa"

**Title**

**Generation of full-sib seed families**

In 2021, we generated full-sib seed families as follows. We sowed 400 seeds in Einheitserde CLASSIC® substrate (PATZER ERDEN GmbH, Sinntal-Altengronau, Germany) and moved them into a phytotron (constant temperature 21°C, 24h light cycle, 60% relative humidity). Two weeks later, we potted each plant into a 7 × 7 cm pot filled with Einheitserde CLASSIC® and moved pots into a greenhouse at the Botanical Garden of the University of Zurich, Switzerland (constant temperature at 21°C day and night, 16h light cycle, 60% relative humidity). From 398 surviving plants, we randomly performed 199 cross-pollinations between non-overlapping pairs of plants. In total, 194 of these pairs produced seeds forming a full-sib seed family from which one we randomly selected 108 full-sib seed families for our experimental evolution (Fig. 1a). These full-sib seed-families are designed to maximize genetic diversity (Dorey et al. 2024; Frachon & Schiestl 2024; Kofler et al. 2024).

**Experimental evolution study design**

Each replicate comprises seeds from those 36 full-sib families, with each family represented by a single randomly selected seed. Our initial generation was composed of 324 plant individuals (3 replicates × 3 treatments × 36 full-sib seed families × 1 seed; Fig. 1b). We randomly allocated the replicate populations to cages, initially and for each generation, under the constraint that replicate populations were not in the same cage position more than once over the course of the entire experiment. We performed this experimental evolution over six generations, from April 2021 to November 2022 (generation 1: April to June 2021, generation 2: July to September 2021, generation 3: September to November 2021, generation 4: April to June 2022, generation 5: June to August 2022, and generation 6: August to November 2022). We watered the plants up to twice a day depending on the weather conditions.

For the Control treatment, we ensured that each plant received pollen from one other plant and donated pollen to one other plant such that each of the 36 plants within a replicate was both donor and recipient of pollen exactly once.

We implemented the pollination treatments in the next three weeks after moving the plants to the outdoor cages. We randomly selected the temporal order of treatment implementation. To avoid pollen transfer from cage to cage (*i.e.*, from replicate population to replicate population), we opened at most one cage on a single day for the Full- and Limited-Access treatments. For the Full- and Limited-Access treatments, we marked open flowers with red tape on the day of the treatment, restricting future fruit counts to flowers that could have been pollinated by pollinators. We also marked the six pollinated open flowers with red tape in Control treatment. Because pollinator activity is lower during cloudy and windy weather (Goodwin et al. 2021; Vicens & Bosch 2000), we performed the Full and Limited Access treatments only on sunny and non-windy days.

**Recording of climate data during experimental evolution**

We recorded temperature in degrees Celsius (°C) and light intensity (in lux) in one-hour intervals over the entire experiment with a battery-driven HOBO^®^ Pendant MX2202 Temperature/Light datalogger installed in the central outdoor cage. For each day of the experiment (over the period from 00:00 am to 11:59 pm time of day), we calculated the minimum, maximum, and mean temperature, as well as the maximum and mean light intensity. We then calculated for each experimental generation the mean and variance of the five temperature and light variables we had recorded across the respective experimental days. We performed a principal component analysis (PCA) using the dudi.pca function of the package ade4 v.1.7-22 (Dray & Dufour 2007) in R environment v4.4.1 (R Core Team 2024) to capture the variation in the ten climatic variables within the two first principal components (PCs).

**Resurrection experiment design**

We specifically resurrected the status from before the experiment by generating three replicate resurrection populations from the three sets of 36 initial full-sib seed families (one offspring per initial family). To resurrect the status from after the experiment, we propagated sixth-generation individuals from each of the nine replicate populations as described below in “Estimation of relative fitness and application of selection”, *i.e.*, with individual contributions proportional to estimated fitness (Fig. 1). We sowed stored seeds at room temperature from the initial populations and from the sixth generation under controlled conditions in a greenhouse as described above (“Generation of full-sib seed families”).

**Characterization of pollinator communities during experimental evolution**

During pollination treatments involving the natural pollinator community of the Botanical Garden of Zürich, we estimated pollinator abundance and diversity that visiting plants. For the Limited Access treatment, we recorded the behaviour of every insect entering the cage and pollinating plants for 20 consecutive minutes. For the Full Access treatment, we recorded pollinator visitation for 20 consecutive minutes every hour for five hours (for a total of 5 × 20 minutes). We categorized pollinators as: “small bee”, “small hoverfly”, “medium bee”, “medium hoverfly”, “honeybee”, “bumblebee”, “*pieris* butterfly”, and “wasp”. We tracked the path of each insect within the cages to determine which plants were visited. For each replicate population within each generation, we calculated the overall *abundance* of insect as the total number of visits recorded. Similarly, we measured the abundance of each category of insects. We estimated the *abundance evenness* as the variance of the distribution of visits across plants. Lastly, we estimated the *Shannon diversity* of pollinators within each replicate population within each generation following this formula:

$$Shannon diversity = - \sum(p_{i} \times ln(p_{i}))$$

with $p_{i}$ the proportion of each category of species.

**Plant traits measured in experimental evolution**

From two days after each pollination treatment, we collected up to three recently opened and unpollinated flowers from each plant to characterize floral morphological traits. We measured the flower diameter with electronic callipers and mounted the petal-less flowers on paper sheets with transparent adhesive tape (Fig. S4). We scanned the sheets with a desktop scanner (imageRunner Advance DX, Canon) with an image resolution of 600 dpi. We used ImageJ software (Schneider et al. 2012) to measure the length and width of the petals (*petal length*, *petal width*), the length of the pistil (*pistil length*), and the length of the longest stamens (*stamen length*). We quantified *herkogamy*, *i.e.*, the spatial distance between anther and the stigma, by calculating the difference between the pistil length and the stamen length. For all traits described above, we calculated the mean per plant as the arithmetic mean across the morphological measurements of the three flowers per plant.

Although *B. rapa* is generally self-incompatible, variation in selfing rates was observed in response to pollinators (Ramos and Schiestl 2019, Kofler et al. 2024). Two days after the pollination treatments, we assessed the ability of plants to self-pollinate by autogamy (pollen fertilizing ovules of the same flower) for each treatment. To assess autogamy, we manually self-pollinated up to three flowers transferring pollen from their own stamens to their stigmas. We marked the flowers pollinated under autogamy with marking tapes of different colours. After seed harvesting (see below for details), we estimated the mean number of seeds under autogamy produced per flower for each plant as the respective arithmetic means. We then calculated the *autogamy ability* by dividing the mean number of seeds under autogamy by the number of fruits self-pollinated.

At fruit maturity, we dried plants in a dry greenhouse (temperature, humidity, and light were not controlled) and we measured plant height. We measured the fruit length for fruits produced from pollination treatments (outcrossing) with an electronic calliper and calculated the *mean fruit length under outcrossing*. For each harvested plant, we counted the total number of seed and calculated the mean number of seed per fruit and per flower for each plant. We weighed the *total number of seeds per plant* with an electronic scale (Mettler Toledo, AG204; precision: 0.001 g) and calculated the *mean seed weight per plant* by dividing the total seed weight by the total number of seeds. We conserved the seeds at room temperature for the next generation.

**Estimation of relative fitness and application of selection**

To propagate plants over six experimental generations and apply selection within treatment populations, we estimated *relative fitness* from the product of the total number of seeds produced times the germination rate. Specifically, after seed harvesting and phenotyping of the first generation, we sowed 10 seeds per parental plant into seed trays (diameter: 5 cm) and kept them in a phytotron for germination (21°C, 24h light, and 60% relative humidity). We estimated the *germination rate,* $g_{i,j}$, of parental plant $i$ in experimental replicate population $j$ by dividing the number of seeds that germinated up to five days after sowing by the total number of seeds sown from that plant. We estimated the relative fitness $w_{i,j}$ of parental plant $i$ in experimental replicate population $j$ as

$$w_{i,j}=\frac{g_{i,j}s_{i,j}}{\sum_{i=1}^{36} g_{i,j}s_{i,j}},$$

where $s_{i,j}$ is the total number of seeds produced by plant $i$ in experimental replicate population $j$. We approximated the expected contribution of each plant to the next generation by rounding half up 36 × $w_{i,j}$ to the closest integer, which be denoted by $\eta_{i,j}=\lceil0.5\lfloor2\times36\times w_{i,j}\rfloor\rceil$. If the sum of all approximated contributions fell above (or below) 36 by *d*, we ranked the parental plants by the rounding difference, $r_{i,j}=\eta_{i,j}-36 w_{i,j}$, and removed (or added) a single offspring from each of the *d* highest (lowest) ranking parental individuals (Fig. 1e). Eight days after sowings, we potted the seedlings in standardized soil as described above. We followed the procedure described above for each new generation during the six generations of experimental evolution.

**Plant traits measured in resurrection experiments**

We phenotyped the second resurrection generation for phenology, floral morphology, and fitness components as follows (Table S1). We recorded the *flowering time* as the number of days after sowing at which the first flower appeared. We measured flower morphology (*mean flower diameter*, *mean petal length*, *mean petal width*, *mean pistil length*, *mean stamen length*, *and mean herkogamy*) for three flowers per plant as described in “Plant traits in experimental evolution” above. In addition, we counted the *total number of secondary inflorescences* (*i.e.*, branches with at least two flowers), the *total number of flowers* per plant, and we measured *plant height*.

To estimate the outcrossing ability of the plants, we collected four stamens from all the individuals of a given replicate resurrection population that we pooled together and mixed with a make-up brush. We applied the mixed pollen to six flowers per plant.

Two days later, we collected floral scent (Ramos & Schiestl 2019). For each plant with at least one recently opened flower, we enclosed one to two full inflorescences in a roasting bag (diam.: 31 cm, Bratschlauch, Toppits). We inserted into the bag a Tenax filter (glass cylinder containing 30 mg of Tenax TA and60/80 mesh from Supelco, Bellefonte, USA) and	pumped 150 mL air per min into the filter for two hours with a self-made electric pump (Schiestl 2014). As a control, we applied the same procedure to a Tenax filter placed into an empty roasting bag positioned in the same room. Immediately after, we wrapped filters into aluminium foil and stored them in a -20 °C freezer. Three to eight weeks later, we extracted	volatiles using gas chromatography with mass selective detection (GC–MSD) as described in Ramos and Schiestl (Ramos & Schiestl 2019). We identified compounds using Agilent MSD ChemStation software (v.E.01.00, Agilent Technologies AG, Santa Clara, USA) by comparing samples mass spectra with the Mass Spectral Library of the National Institute of Standards and Technology (NIST) database and synthetic standards of all compounds. We quantified compounds by converting each peak area into absolute amounts using calibration curves derived from synthetic standards. For down-stream analyses, we only considered compounds that were absorbed by the test filters but not by the control filter.

At fruit maturity (*i.e.*, about three weeks after hand-pollination), we dried plants into the greenhouse as described in ’Plant traits measured in experimental evolution’ above. We measured *mean fruit length under outcrossing* with electronic callipers, and counted the *number of seeds per fruit under outcrossing* for all fruits per plant. For flower morphology traits, fruit lengths, and the number of seeds per fruit, we calculated the arithmetic mean per plant for each trait. We weighed outcrossed pollinated seeds with an electronic scale (Mettler Toledo, AG204; precision: 0.001 g) and we calculated the *mean* *seed weight* per plant and the *mean seed weight per fruit*.

To assess plants’ ability to outcross and to self-pollinate via autogamy, we hand-pollinated plants from the ‘third’ resurrection generation. To test outcrossing ability, we pollinated within each replicate resurrection population three flowers of each plant with pollen collected from all individuals in the respective replicate resurrection population. To test the ability to self-pollinate by autogamy, we hand-pollinated three flowers of each plant with pollen from the respective plant. At fruit maturity, we measured the length of fruits from outcrossing and selfing and counted the seed number per plant. We calculated the *mean fruit length under outcrossing*, the *mean fruit length under autogamy*, the *mean number of seeds under outcrossing,* and the *mean number of seeds under autogamy* per plant.

**Additional statistical analyses**

We checked normality of the 17 traits using Shapiro tests in R v4.4.1 (R Core Team 2024) for each replicate population. For statistical analyses requiring normality, we discarded traits that failed the Shapiro test (*p* value < 0.01) and could not be normalized through transformation (boxcox, square root, log(1+x), and x^2^) (n = 7 traits discarded).

To verify if the overall abundance, the abundance evenness, and the Shannon diversity of pollinators statistically differed between the Limited and the Full Access treatments, we analysed each of the three ecological factors separately with an Anova in R with the following formula:

$$\begin{matrix} Abundance \\ Abundance evenness \\ Shannon diversity \end{matrix} \sim treatment +climatePCs + \begin{matrix} Number of \\ flowers \end{matrix}+ \begin{matrix} Days since \\ sowing \end{matrix}$$

where *treatment* represents the three pollination treatments (fixed factor), *climatePCs* the first two principal components derived from the climate PCA (fixed factor), *Number of flowers* the total number of open flowers in the cage on the day of the pollination treatment (fixed factor), and *Days since sowing* the number of days between the sowing and the pollination treatment (random factor).

**Additional information regarding the four-choice test of pollinator preference**

We randomly picked each set of four plants from within the respective replicate population under the constraint that we used each individual plant at most once. We conducted the four-choice tests in two flight cages (1.75 m × 1.8 m × 1.2 m; ca. 240 mesh per square inch, Spatz Zelte & Reparaturen AG) under the same temperature and light conditions as those applied during resurrection experiment.

We conducted the four-choice tests between 9:00 am and 4:00 pm over three consecutive days. We used bumblebee (*Bombus terrestris*) and hoverfly (*Episyrphus balteatus*) as models for insect pollinators, both of which visited *B. rapa* flowers in our common-garden experiment. *B. terrestris* hives and *E. balteatus* pupae were brought from Andermatt Biocontrol Suisse AG (Grossdietwil, Switzerland). We released a given pollinator individual at most once to avoid a confounding effect from learning or habituation processes.

We unravelled which phenotypic traits were involved in plant attractiveness to pollinators. To include dependency among the four focal plants used in a same four-choice test, we performed a multinomial logistic regression with the mlogit R package (Croissant 2020). For all phenotypic traits simultaneously, we explained the bumblebee and the hoverfly choices by all phenotypic traits as explanatory variables. Following the mlogit package’s syntax, we included the identification number of each four-choice test as “choice index” and the treatment as “alternative index”. To include all confounding effects that could influence the analysis, we added the position of the pots (“top-left”, “top-right”, “bottom-left”, and “bottom-right”) as “alternative specific” variable. Pollinator species, replicate, day of the four-choice tests, and the identification of the cage which was used (“left” or “right”) were defined as “choice specific” variables. In addition to those models, we performed another set of multinomial logistic regression with all the same parameters, but for bumblebee and hoverfly choices separately and without the pollinator species as an explanatory variable. Those additional models allowed us to decipher the bumblebee or hoverfly personal preference from their shared preference. The replicate, the identity of the cage, beta-pinene, benzaldehyde, and plant height have been later discarded from the models for never being significant. Butene isothiocyanate was also discarded for being too correlated to limonene (Spearman rho = 0.87***).

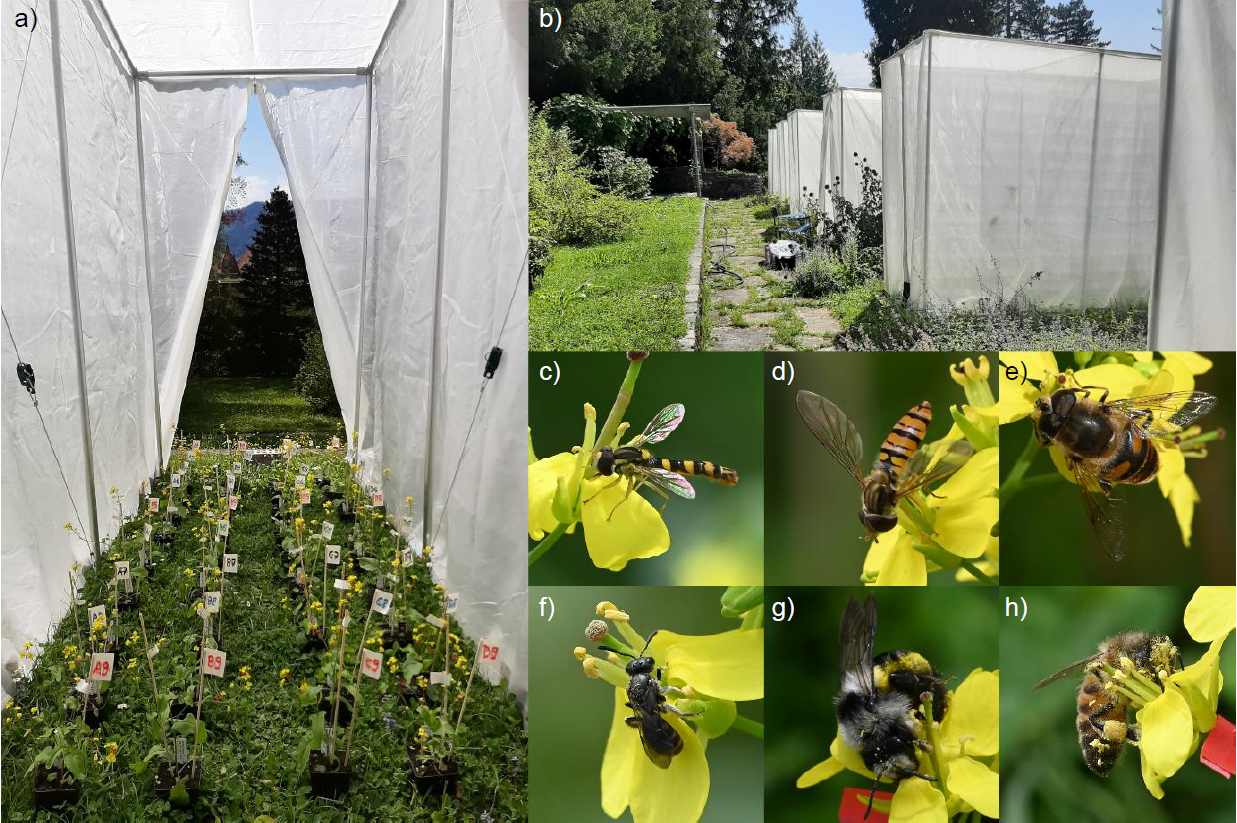

**FIGURE S1 |** **Outside setting of the experimental evolution**. a) Inside and b) outside view of one of the nine cages during pollination treatment of experimental evolution. Observations of c) *Sphaerophoria scripta*, d) *Episyrphus balteatus*, e) *Eristalis tenax*, f) *Chelostoma distinctum*, g) *Bombus pratorum*, and h) *Apis mellifera* pollinating flowers of *B. rapa* inside the cages during the six generations of the experimental evolution.

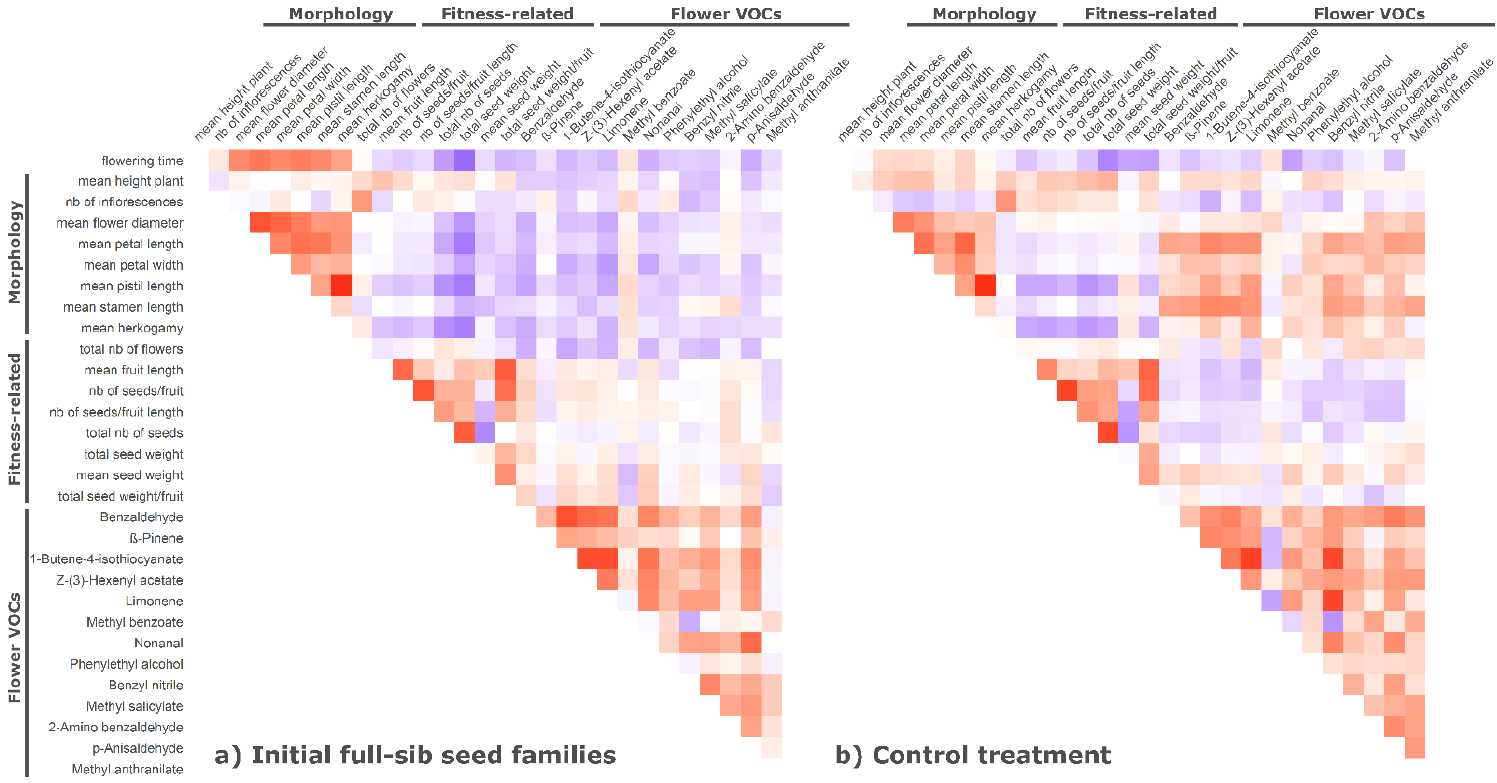

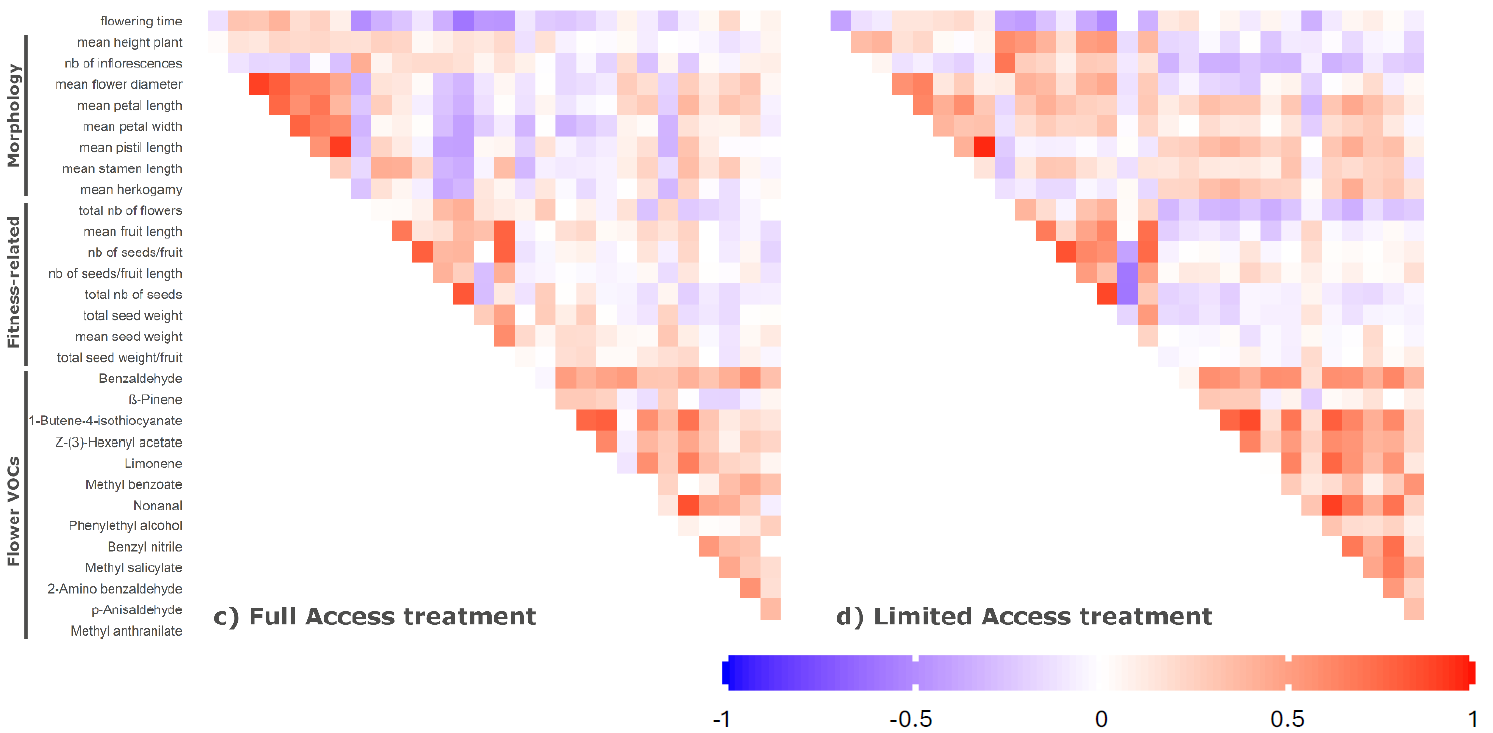

**FIGURE S2 |** **Pairwise Spearman correlations among traits within the different populations**. Phenotypic traits (Morphology and floral scent, fitness components and compound fitness proxies have been measured in the resurrection experiment for a) the initial sib-seed families, b) the Control, c) the Full Access, and d) the Limited Access treatments separately. Statistically significant (p < 0.05) Spearman’s rank correlation coefficients are shown as filled squares. The intensity and gradient of the colour are proportional to the correlation coefficient.

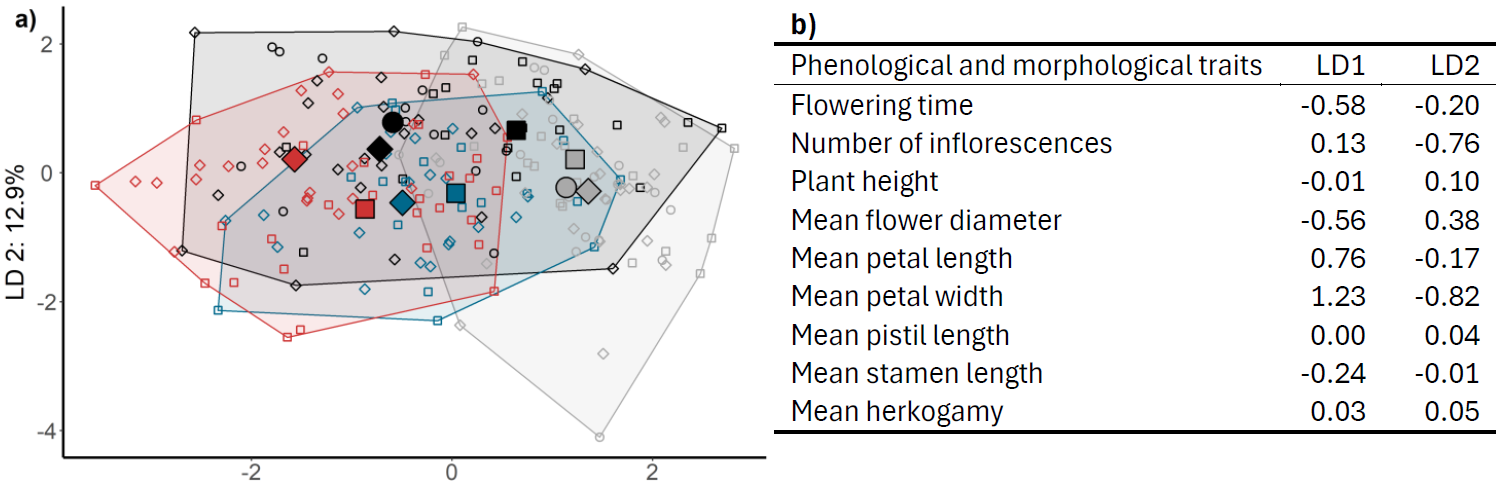

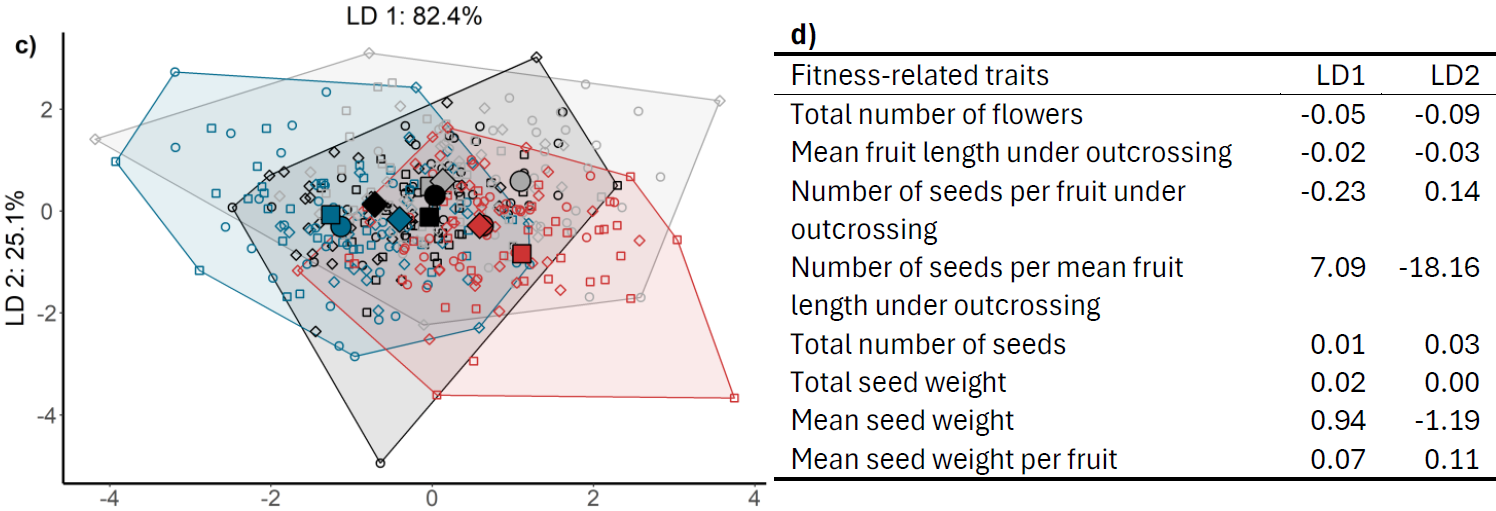

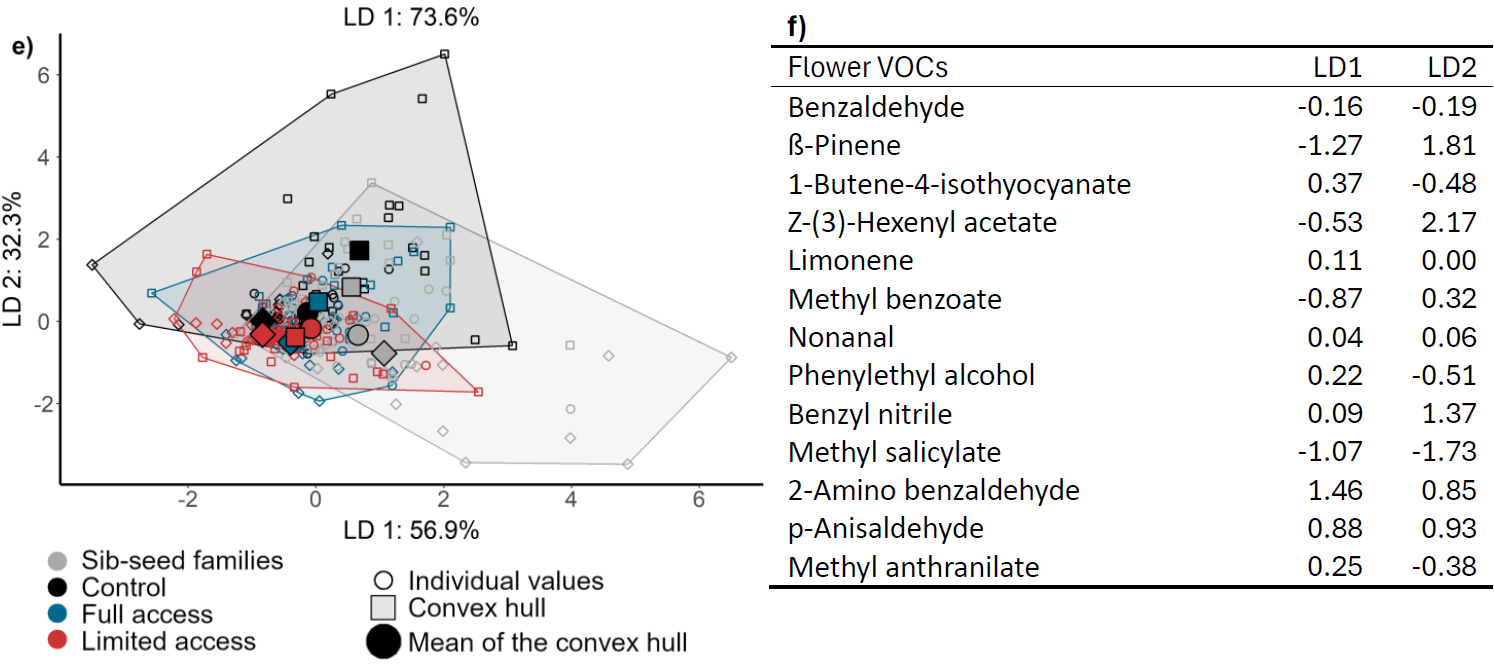

**FIGURE S3 |** **Phenotypic differentiation among the initial full-sib seed families and the three groups of populations evolved under different pollination treatments revealed in the resurrection experiment**. Phenotypic differentiation was quantified in a linear discriminant (LD) analysis for a) and b) phenological and morphological traits, c) and d) fitness components and fitness proxies, and e) and f) floral VOCs. Means across replicate populations are shown along LD axis 1 and 2, and are coloured by their respective pollination treatment. The two LD axes are annotated with linear combinations of phenotypic traits that aligned most closely with the respective LD axis. b), d), and f) show the values of the scores obtained for each trait for the two LDs of respectively the a), c), and e) LDA.

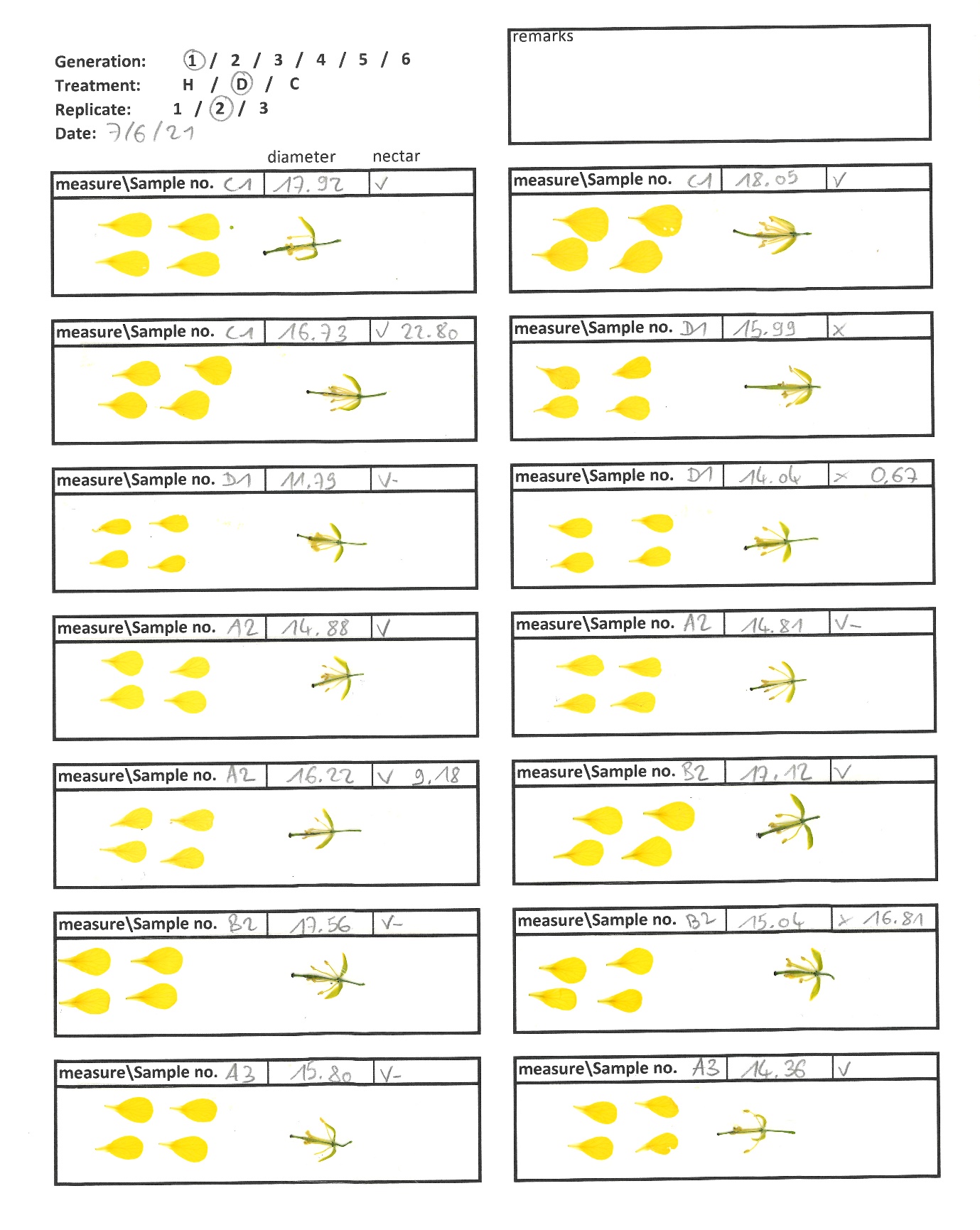

**FIGURE S4 |** **Example of a data sheet with mounted flowers**. Flowers have first been dissected and their petals and reproductive organs mounted separately with transparent adhesive tape.

**TABLE S1 |** List of all phenotypic and fitness-related traits measured. The two columns on the right of the table specify which traits were measured in the experimental evolution study, the resurrection approach, or both. The names of traits used in this paper, their description, and their metrics are indicated here.

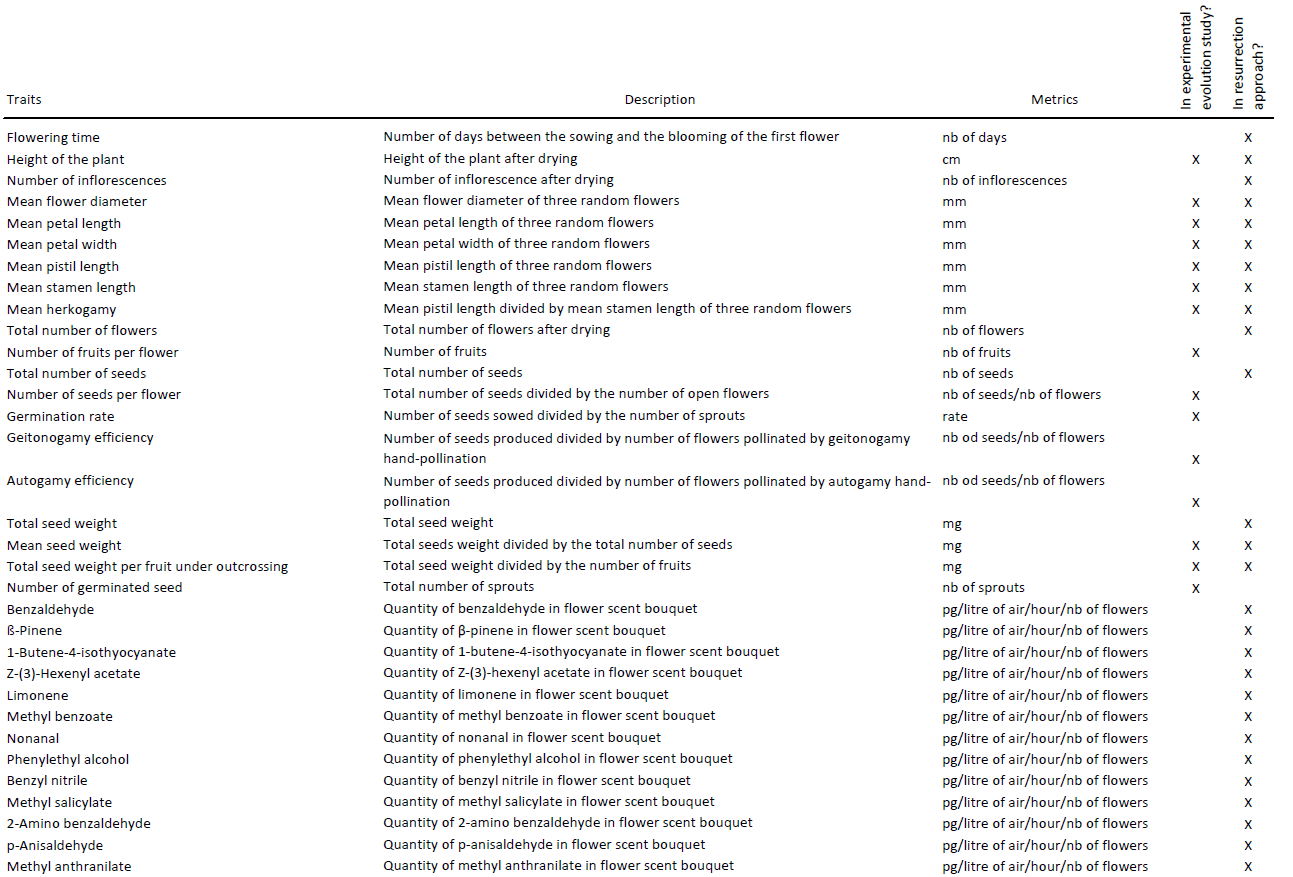

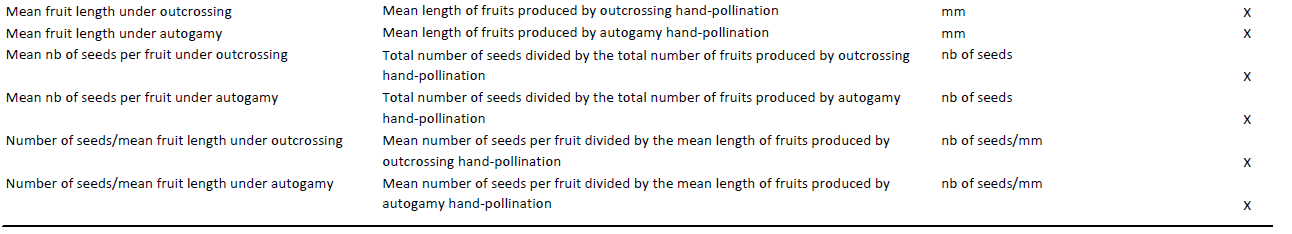

**TABLE S2 |** Contribution of each climatic variable on the five first principal components. Values in bold indicate for each variable the highest contribution in the first and second components.

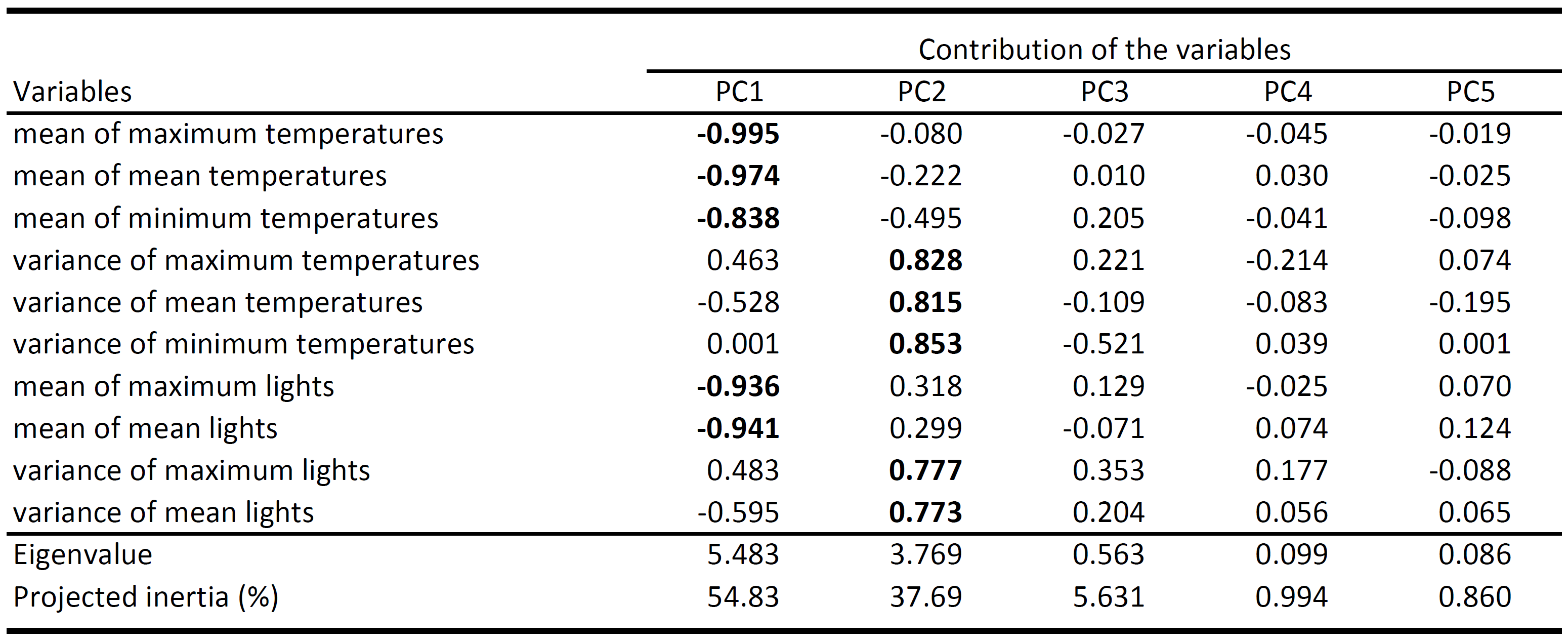

**TABLE S3 |** Variation of insect abundance and diversity between Limited and Full Access treatments during the experimental evolution study. The three ecological indexes have been explained by pollination treatment (fixed effect), total number of open flowers in the cage on the day of the treatment (fixed effect), two climate variables (fixed effects), and the number of days between the sowing of the seeds and the day of the treatment (random effect). The Full access treatment is included within the residuals. The coefficient of the Treatment effects represents then the effect of the Limited access treatment. Bold variables indicate explanatory variables with *p*-values inferior to 0.05.

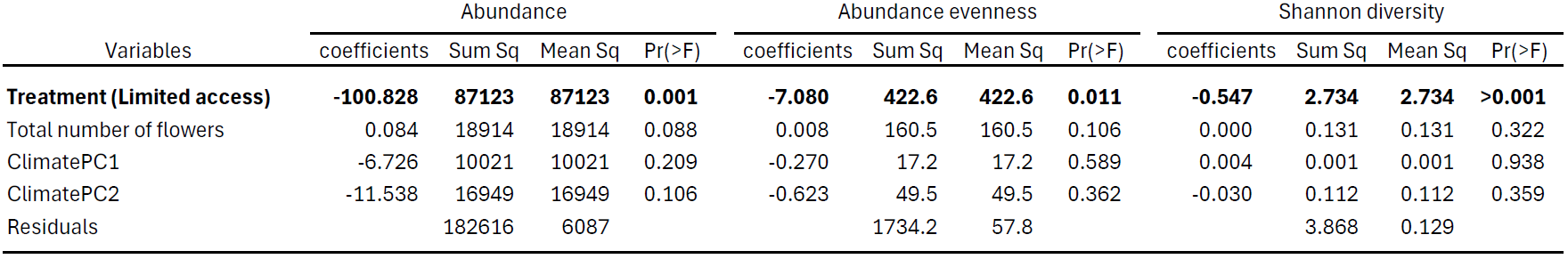

**TABLE S4 |** Results of pairwise Kruskal—Wallis tests performed on 17 measured traits measured during the experimental evolution. Two Kruskal—Wallis tests have been performed on: (1) overall differences among treatments (treatments), and (2) overall differences across the six generations (generation). *p*-value are indicated in the table with significant ones (*p*-value < 0.01) in bold.

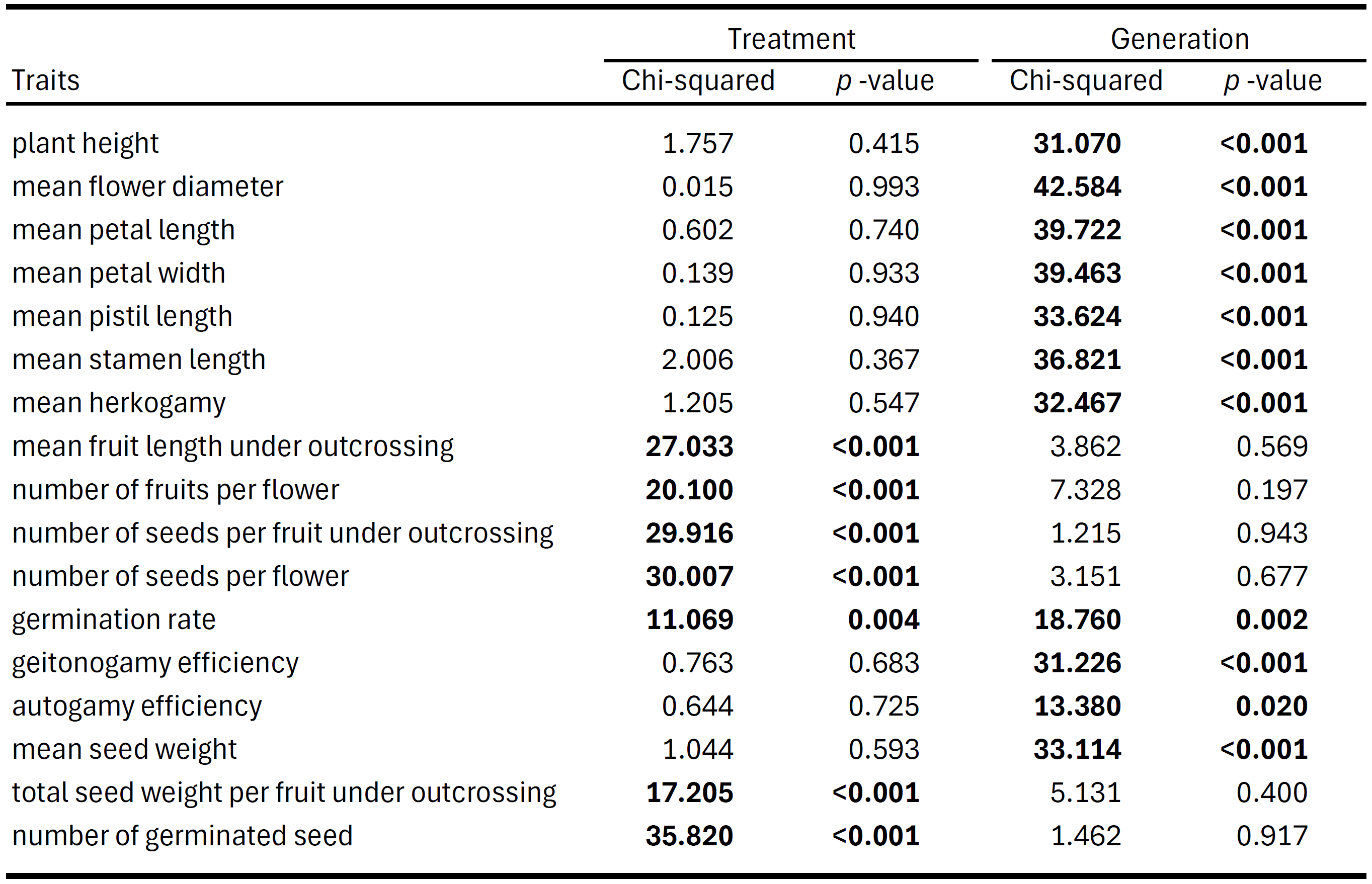

**TABLE S5 |** Directional selection on phenotypic traits within different resurrection treatments. Linear selection gradients (directional selection) have been obtained within each treatment separately (including the initial sib-seed families), by explaining the estimated relative fitness with phenotypic traits in a linear model. All phenotypic traits have been standardized prior to the analysis. a) With all plant architecture and flower morphology traits, and b) with all flower VOCs. Traits highly correlated to another (corr > 0.8 or corr < -0.8) have been excluded. Bold variables indicate explanatory variables with *p*-values inferior to 0.05.

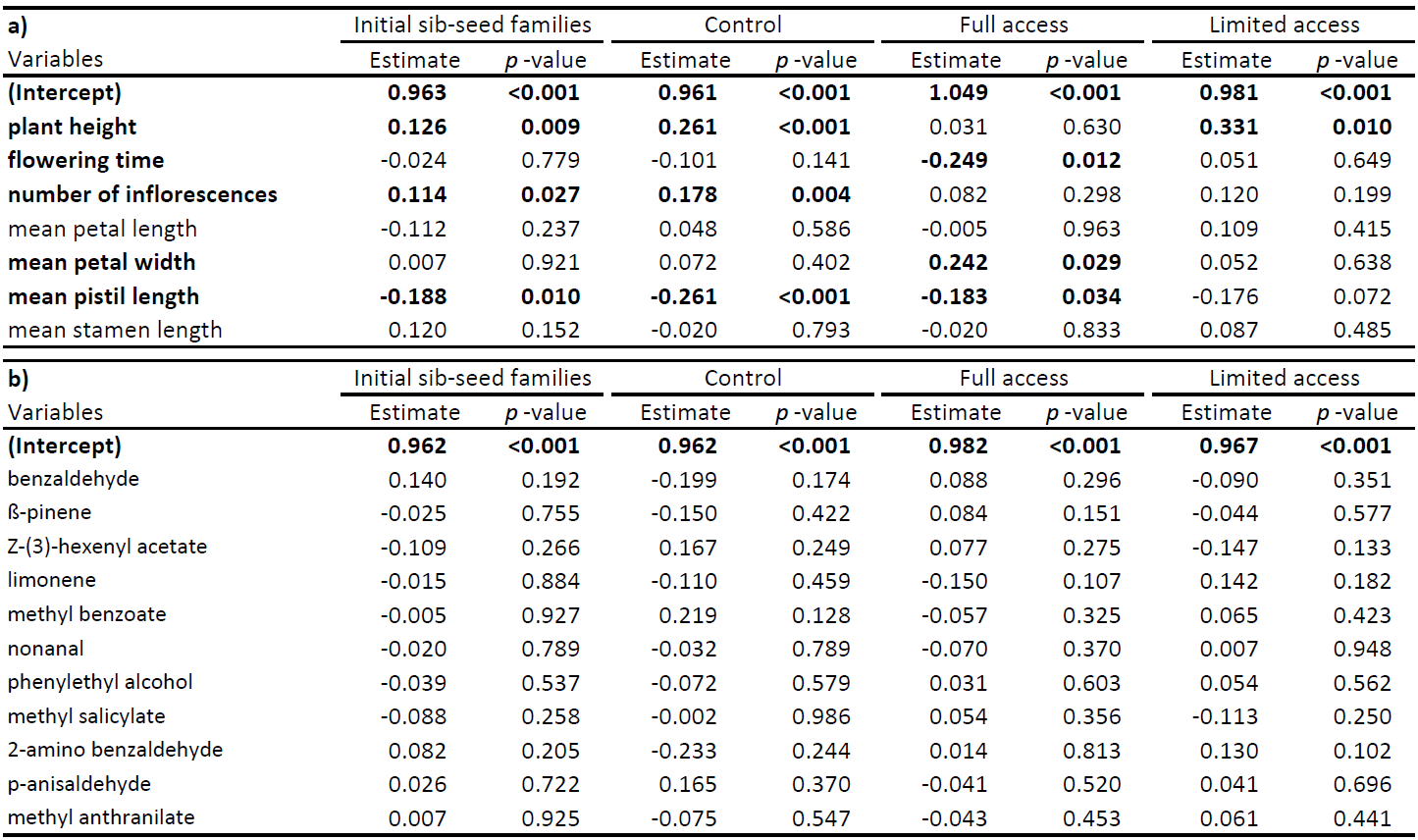

**TABLE S6 |** Insect preference on 22 phenotypic traits. Results of Wilcoxon test on phenotypic traits between plants chosen and not chosen by pollinators. For each trait, the arithmetic means between plants chosen and not chosen by either bumblebees or hoverflies is indicated. Bold variables indicate explanatory variables with p-values inferior to 0.05.

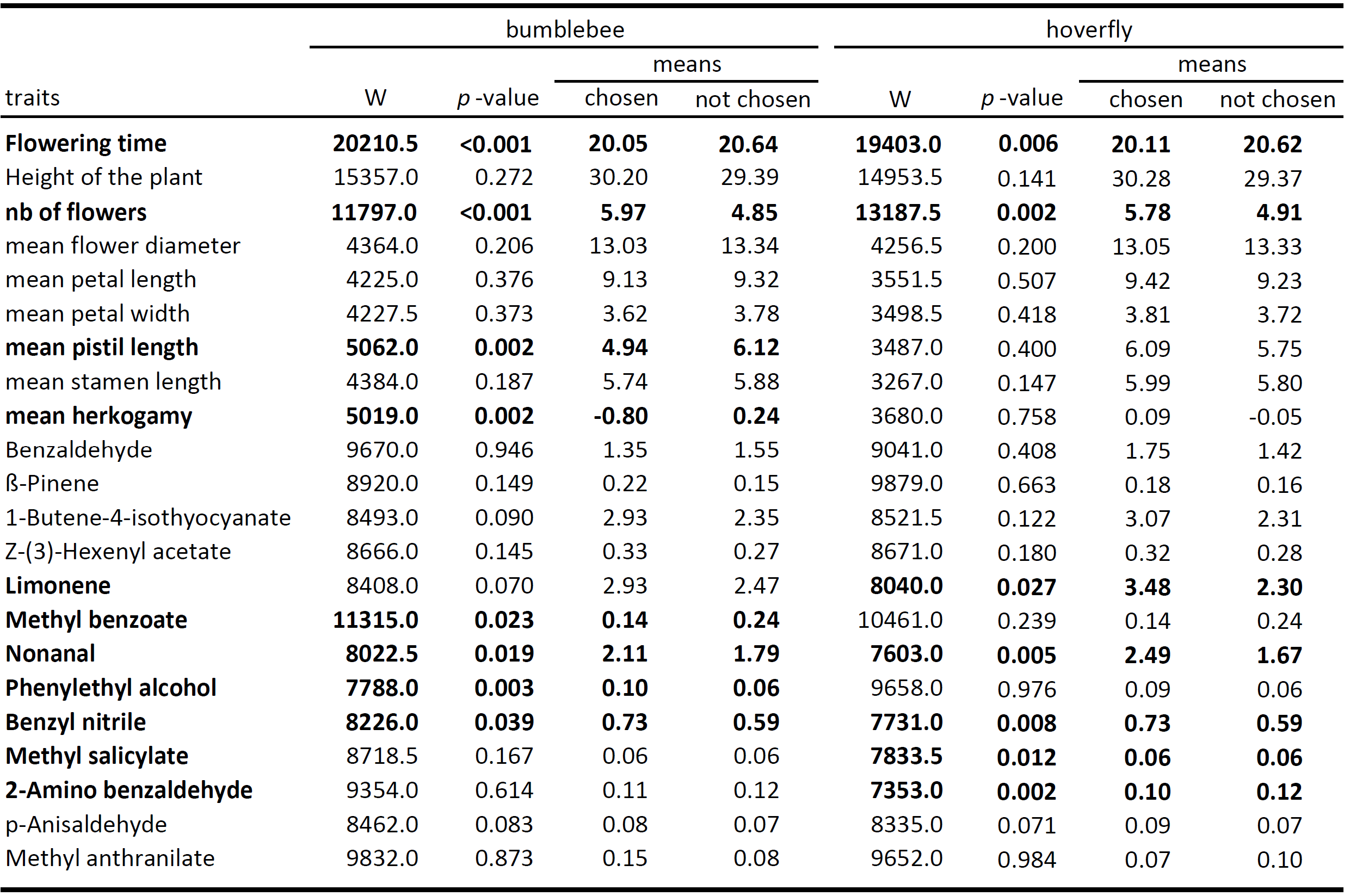

**TABLE S7 |** Insect preference on phenotypic traits. Results of multinomial logistic regression: pollinator choice has been explained with treatments, phenotypic traits, replicate, position of the pots in the cage, the cage position, and date (day of the four-choice tests). Every statistical analysis has been performed separately between four-choice tests with bumblebees and hoverflies.

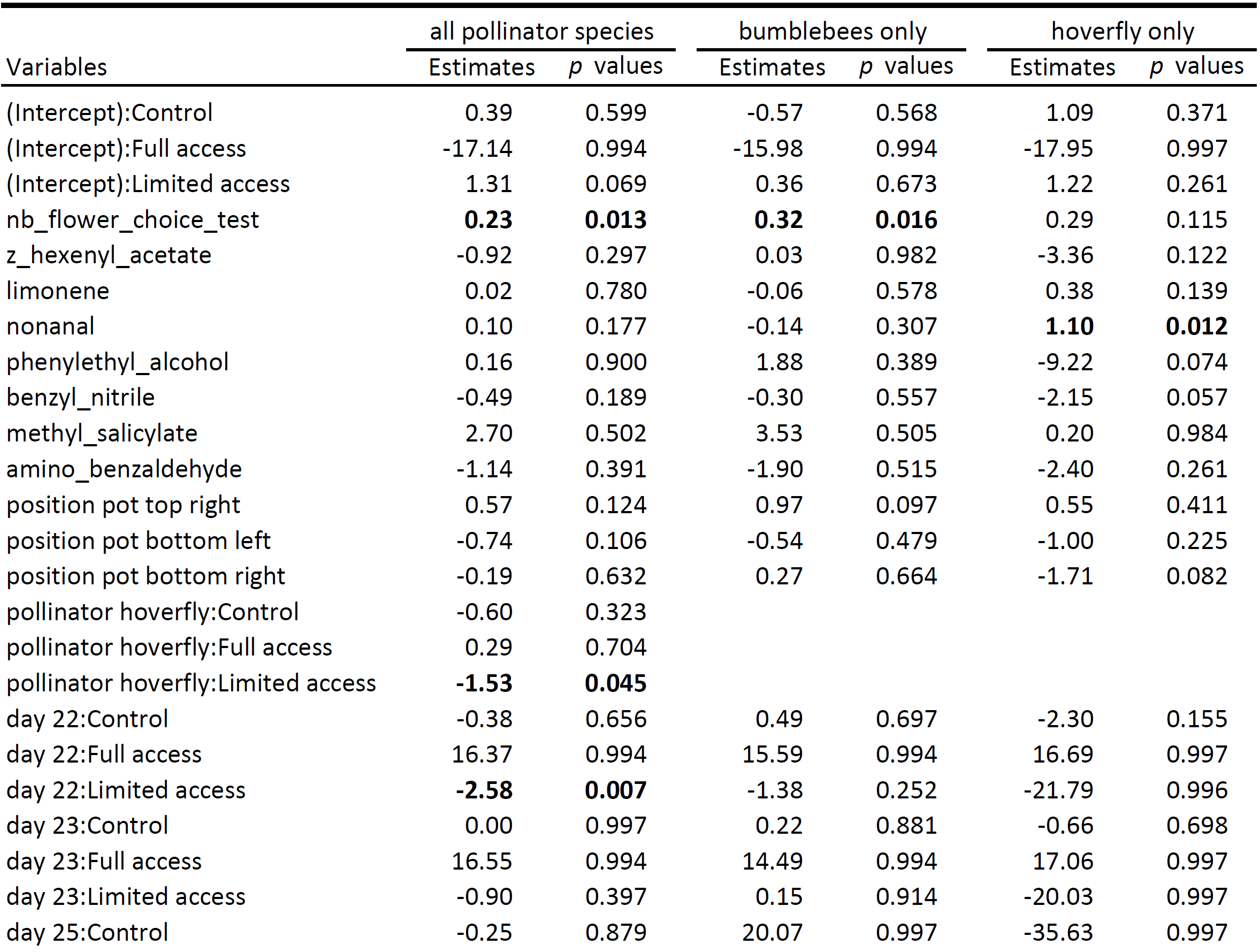

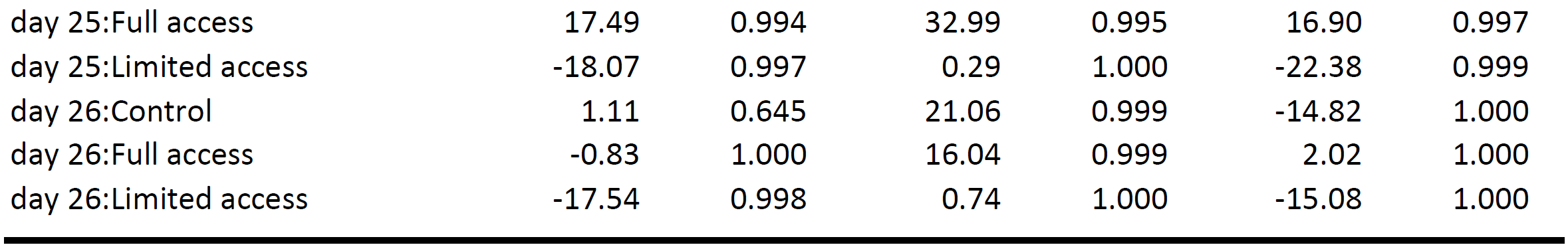
